# 4-Hydroxynonenal suppresses IL-10 production during infection

**DOI:** 10.1101/2025.11.24.690094

**Authors:** Melina Ioannidis, Sjors Maassen, Lisanne Boekhoud, Martijn den Ouden, Mihai Simioniuc, Pieter Grijpstra, Danny Incarnato, Frans Bianchi, Hjalmar Bouma, Geert van den Bogaart

## Abstract

Sepsis is a syndrome of life-threatening multiple organ failure induced by infection and hallmarked by the increased production of inflammatory cytokines and reactive oxygen species (ROS). The oxidation of lipids by ROS produces 4-hydroxynonenal (4-HNE), a highly reactive aldehyde that forms adducts with proteins and thereby impacts immune signaling. In this study, using blood samples from patients with sepsis at the emergency department, collected by the Acutelines data- and biobank, we show that 4-HNE selectively suppresses the production of the anti-inflammatory cytokine interleukin (IL)-10, while pro-inflammatory IL-6 and tumor necrosis factor (TNF)-α are unaffected. Mechanistically, 4-HNE causes a pronounced transcriptional reorganization, leading to metabolic reprogramming and activation of HIF-1α signaling. In turn, this suppresses IL-10 production through inhibition of nuclear factor kappa-light-chain-enhancer of activated B-cells (NF-κB) signaling, whereas IL-6 and TNF-α are unaffected due to increased activation of p38 mitogen-activated protein kinase (MAPK) signaling. This suppression likely occurs in sepsis, because, whereas overall 4-HNE protein adduct levels are increased in blood samples of sepsis patients, they are decreased in monocytes and T cells and negatively correlate with IL-10 levels. Thus, our data show that 4-HNE selectively suppresses IL-10 production in sepsis. This is likely relevant to the clinical outcome of sepsis patients because IL-10 levels correlate with mortality.

**Graphical abstract:** 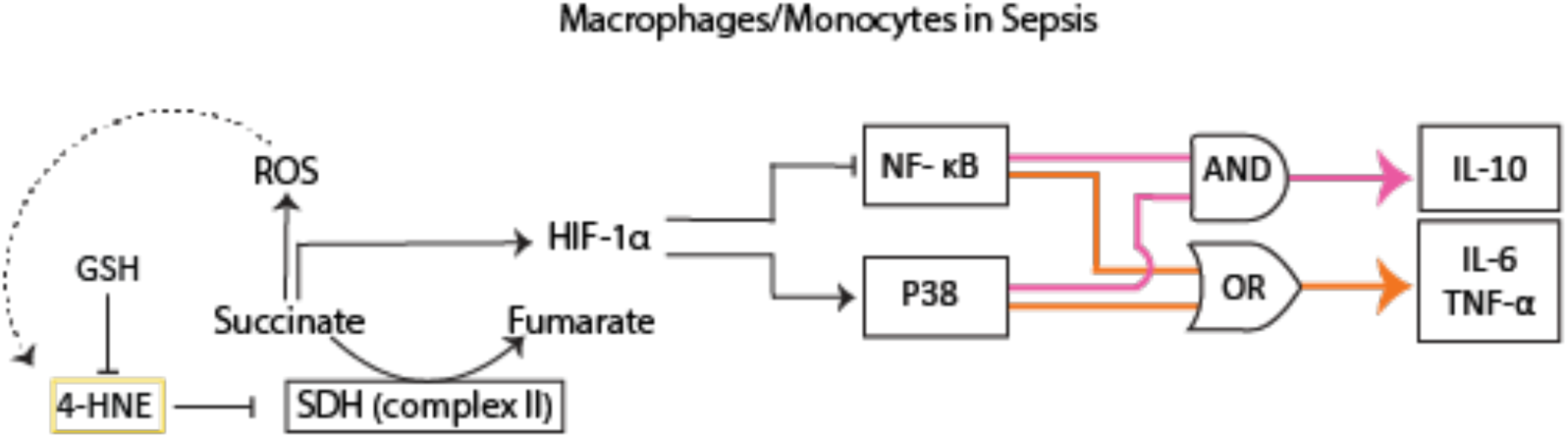

## Introduction

Sepsis is a life-threatening syndrome that arises when the immune response to an infection causes uncontrolled systemic inflammation, which in turn can lead to multiple organ failure (*1*). Sepsis is one of the leading causes of death worldwide, responsible for 11 million deaths in 2017. Half of all 48.9 million global annual sepsis cases occur among children, with 2.9 million global deaths in children under the age of 5 years (*2*). Current sepsis treatment is limited to source control (i.e., administration of antimicrobials and removal of infectious material) and supportive care (e.g., intravenous fluids, vasopressor drugs, ventilation). Since the exact mechanisms leading to organ failure in sepsis remain unknown, targeted therapy directed at the root cause of organ failure is not available, and the morbidity and mortality rates remain high (*3*, *4*).

In sepsis, immune cells, such as macrophages, monocytes, and T cells, produce increased levels of pro-inflammatory cytokines, including tumor necrosis factor-alpha (TNF-α) and interleukin-6 (IL-6). Consecutively, immune cells simultaneously produce anti-inflammatory IL-10, potentially to limit the inflammatory response (*5*).

The pro-inflammatory systemic inflammatory response syndrome (SIRS) was previously believed to be followed by a compensatory anti-inflammatory response syndrome (CARS). However, nowadays, it is clear that both SIRS and CARS can occur at the same moment but to different degrees, which is patient-dependent, requiring precision diagnostic and tailored treatment (Simona Mera et al., 2011; Ward et al., 2008). Moreover, the role of anti-inflammatory IL-10 in sepsis is complex. In mice, CD169^+^ macrophages control inflammation via IL-10, as macrophage deletion of IL-10 increased lethality upon lipopolysaccharide (LPS) injection, a model for sepsis, and this could be rescued by the addition of recombinant IL-10 (*8*). In line with these findings, the early administration of IL-10 neutralizing antibodies in the cecal ligation and puncture (CLP) mouse model for sepsis aggravated mortality (*9*, *10*). However, the inhibition of IL-10 at 12h after inducing CLP increased survival (*9*), showing that IL-10 can also be detrimental. Evidence suggests that IL-10 can have detrimental effects in human sepsis as well. First, increased plasma levels of IL-10 in 11 patients with recent onset of septic shock correlated with a greater degree of organ failure (*11*). Similarly, in another cohort of 20 sepsis patients, an increase in IL-10 mRNA in peripheral leukocytes predicted poor disease outcome (*12*). Thus, IL-10 is an essential modulator in sepsis but can have both protective and detrimental effects.

In addition to cytokines, macrophages and other immune cell types produce large amounts of reactive oxygen species (ROS) upon pathogen encounter (*13*). The primary sources of cellular ROS are NADPH oxidases in the cytoplasm and plasma membrane, and oxidative respiration in mitochondria (*14*, *15*). ROS kill invading pathogens and are important immune signaling molecules (*14*, *16*). However, excessive amounts of ROS cause oxidative modifications of proteins and other biomolecules and contribute to the uncontrolled immune response in sepsis (*17*, *18*). For example, ROS can oxidize omega-6-polyunsaturated fatty acids (PUFA), producing reactive aldehydes. The most abundant reactive aldehyde produced by lipid oxidation is 4-HNE (*19*, *20*).

4-HNE is a reactive α-β-aldehyde that contains three reactive groups: an aldehyde at C1, a hydroxyl group at C4, and a double bond between C2 and C3. Hence, 4-HNE can form adducts with histidine, lysine, and cysteine residues of proteins and alter their functions (*21*, *22*). Under homeostatic conditions, the concentration of 4-HNE is between 0.05 μM and 15 μM in human blood and serum (*23*, *24*). However, during oxidative stress, the concentration of 4-HNE can rise to more than 100 μM, which has a toxic effect on cells and can trigger apoptosis (*25*). To prevent this, cells can reduce 4-HNE through alcohol dehydrogenase and aldose reductase, oxidize 4-HNE through aldehyde dehydrogenase, or conjugate it to glutathione through a spontaneous or glutathione-S-transferase (GSH)-catalyzed reaction (*26*, *27*).

4-HNE adduct formation alters immune functions. For example, elevated concentrations of 4-HNE (25 μM to 50 μM) stimulate an increase in cyclo-oxygenase (COX) -2 mRNA and trigger the activation of the p38 MAPK signaling pathway in RAW264.7 murine macrophages (*28*, *29*). Within the human monocytic cell line THP-1, 4-HNE dose-dependently inhibits NF-κB activation by preventing the phosphorylation of the inhibitory protein IκBα (*30*).

In this study, we investigate the role of 4-HNE in sepsis. Considering the increased ROS production in sepsis patients (*31*, *32*), we hypothesized that the severity of sepsis correlates with 4-HNE levels. Moreover, we hypothesized that 4-HNE influences sepsis outcomes by altering immune signaling. To test these hypotheses, we investigated the levels of 4-HNE protein adducts in CD14^+^ monocytes and CD3^+^ T cells in buffy coats obtained from patients at the emergency department with either sepsis with a severe outcome (i.e., ICU admission, death, or a rise in SOFA-score of two or more within 72 h), sepsis without a severe outcome, and a control group without sepsis. These 4-HNE adduct levels were correlated with plasma cytokine levels. In parallel, we assessed the effects of 4-HNE on cultured CD14^+^ monocyte-derived macrophages stimulated with the canonical bacterial stimulus LPS.

## Results

To determine the 4-HNE protein adduct levels, we selected buffy coats and serum samples from the Acutelines biobank from three groups of patients (Table 1). The groups were selected to ensure comparability in gender and age distribution. Patients in sepsis groups have confirmed bacterial infections, and for one-third of the patients in both severe and non-severe outcomes, there is a concurrent viral infection. Sepsis severe outcome is defined as patients diagnosed with sepsis who were admitted to the ICU and died within 72 hours of admission.

**Table 1:**
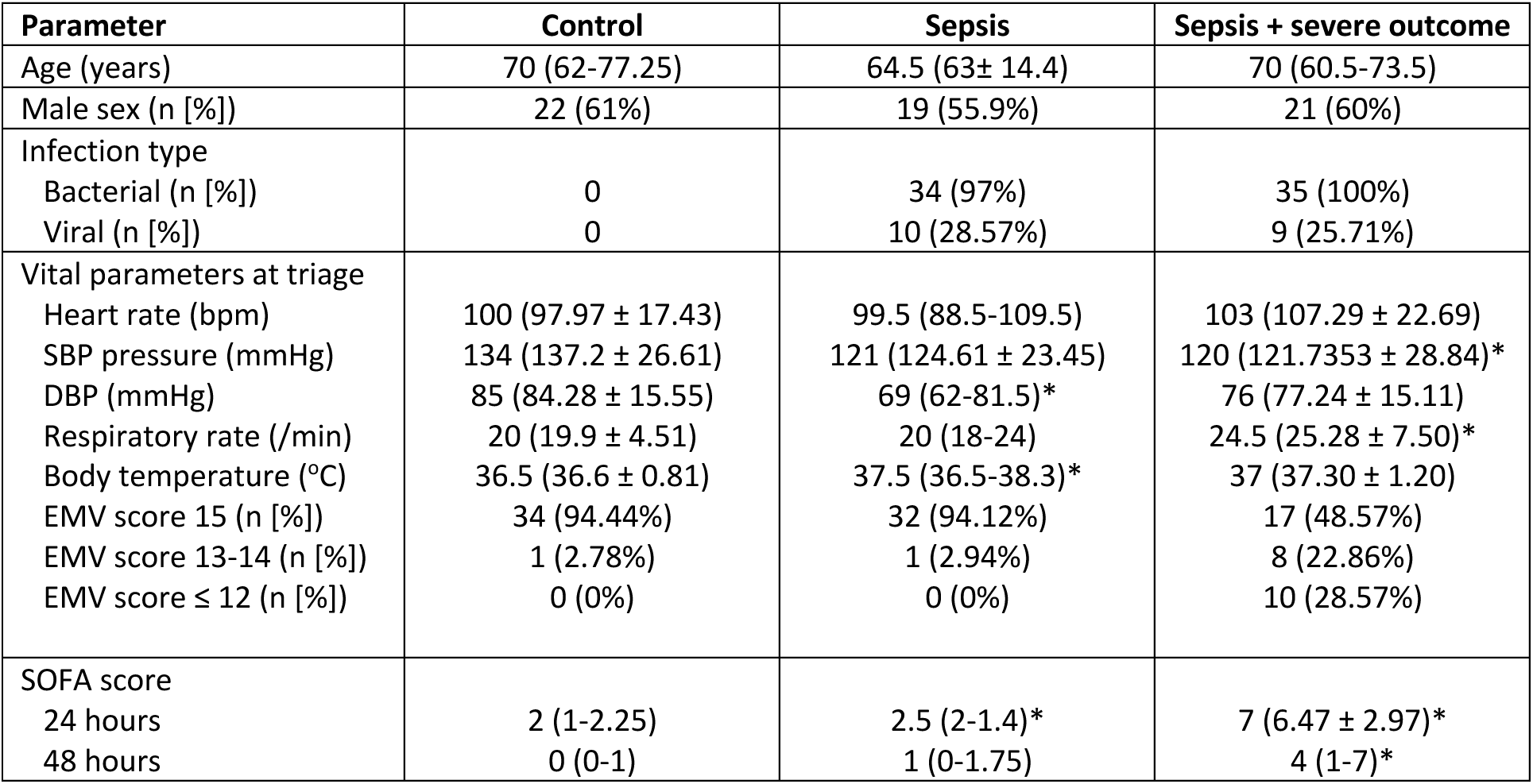
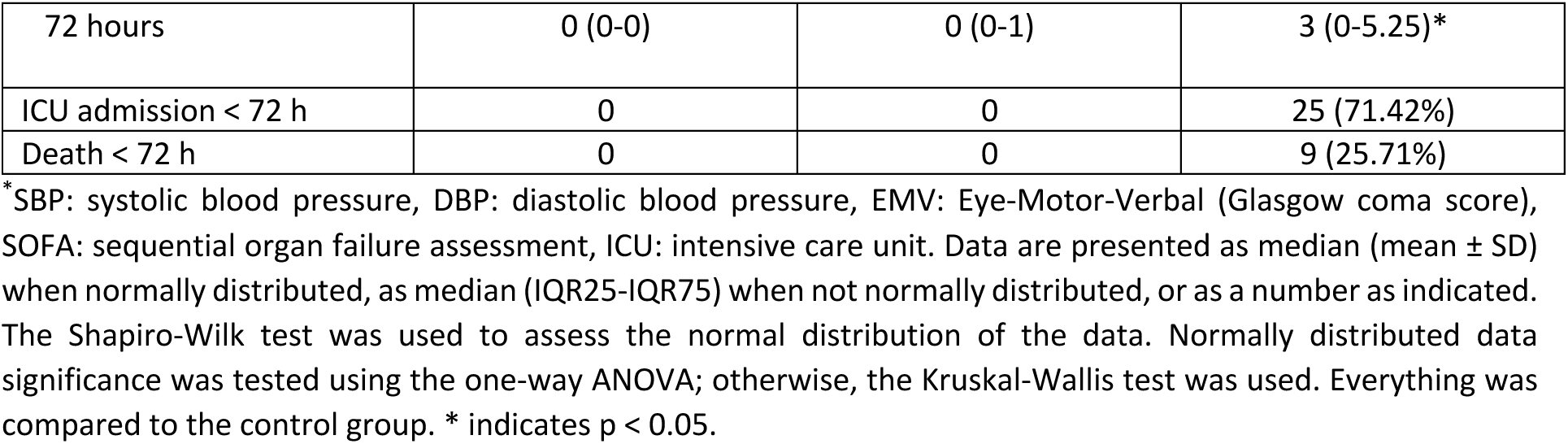
Patient characteristics and clinical outcomes*.

A compensated vasodilated state is commonly observed in sepsis patients, often resulting in a significant decrease in diastolic blood pressure (DBP) while systolic blood pressure (SBP) remains stable. This state can act as a warning sign of impending cardiovascular collapse (*33*). In this study, detailed clinical assessments showed that DBP was notably lower in the sepsis group compared to the control group, while SBP remained unchanged. Moreover, the body temperature in sepsis with non-severe and severe outcomes was significantly increased. Septic patients with severe outcomes displayed a higher decrease in consciousness compared to septic patients without severe outcomes.

The Sequential Organ Failure Assessment (SOFA) score is a widely used tool to evaluate the severity of organ dysfunction in critically ill patients, particularly those with sepsis. It quantifies the degree of dysfunction across multiple organ systems, with higher scores indicating more severe organ failure and an increased mortality risk (*34*). In this study, it was observed that both severe and non-severe sepsis outcomes initially had a SOFA score higher than 2. However, in non-severe cases, the score was reduced to 0, while in severe cases, it remained above 2 even 72 hours after admission to the ED.

The hematological parameters (Table 2) show that C-reactive protein (CRP), IL-6, and IL-10 increased significantly in both sepsis groups, with the highest increase in the severe outcome group.

**Table 2:** Hematological and biochemical parameters upon triage at the ED*.

| Parameter | Control | Sepsis | Sepsis + severe outcome |
| --- | --- | --- | --- |
| Leukocyte count ($\times 10^9/L$) | 10.4 (9.83 ± 4.62) | 12.55 (8.65-16.05) | 12.5 (12.98 ± 7.97) |
| Plasma bilirubin ($\mu\text{mol/L}$) | 8.5 (8.19 ± 3.29) | 11 (6.5-15.5) | 10.5 (6.25-21.5) |
| Plasma creatinine ($\mu\text{mol/L}$) | 94 (71.75-136.5) | 98 (66.25-183.75) | 122 (84-217) |
| Plasma CRP (mg/dL) | 14.5 (3.7-33.5) | 129.5 (52.25-202.25)* | 67 (32.5-187.5)* |
| Serum IL-6 (pg/mL) | 48.35 (41.40-63.57) | 125.72 (70.13-321.42)* | 253.86 (84.8-2037.05)* |
| Serum IL-10 (pg/mL) | 22.77 (14.69-37.11) | 41.53 (21.35-68.57)* | 74.65 (35.95-146.71)* |
| Serum TNF-α (pg/mL) | 392.68 (240.05-481.26) | 312.97 (226.34-439.92) | 291.49 (230.16-419.45) |

\*Data are presented as median (mean ± SD) when normally distributed or as median (IQR25-IQR75) when not normally distributed. The Shapiro-Wilk test was used to assess the normal distribution of the data. Normally distributed data significance was tested using the one-way ANOVA; otherwise, the Kruskal-Wallis test was used. Everything was compared to the control group. \* indicates $p < 0.05$ .

### 4-HNE adducts are increased in patients with sepsis but decreased in monocytes and T cells

For measuring 4-HNE protein adduct levels, we first tested our method on buffy coats isolated from healthy donors that were frozen down immediately after drawing blood. This procedure mimics the buffy coat collection procedure of the Acutelines biobank. The buffy coat from the healthy donor was treated with 10 μM or 50 μM of 4-HNE for 30 min before being stained with an antibody against 4-HNE protein adducts. 4-HNE protein adducts increased with an increased dose of 4-HNE as measured by flow cytometry (Figure 1A). Next, we measured 4-HNE protein adducts in buffy coats from the sepsis patients. As expected, patients with sepsis without or with a severe outcome (i.e., ICU admission or death < 72 hours) have increased levels of 4-HNE protein adducts in the buffy coat as compared to acutely ill patients without an infection that served as the control group (Figure 1B). These results are in line with the increase in malondialdehyde (MDA), which is another lipid peroxidation product, observed previously in the plasma of sepsis patients (*35*).

**Figure 1:**
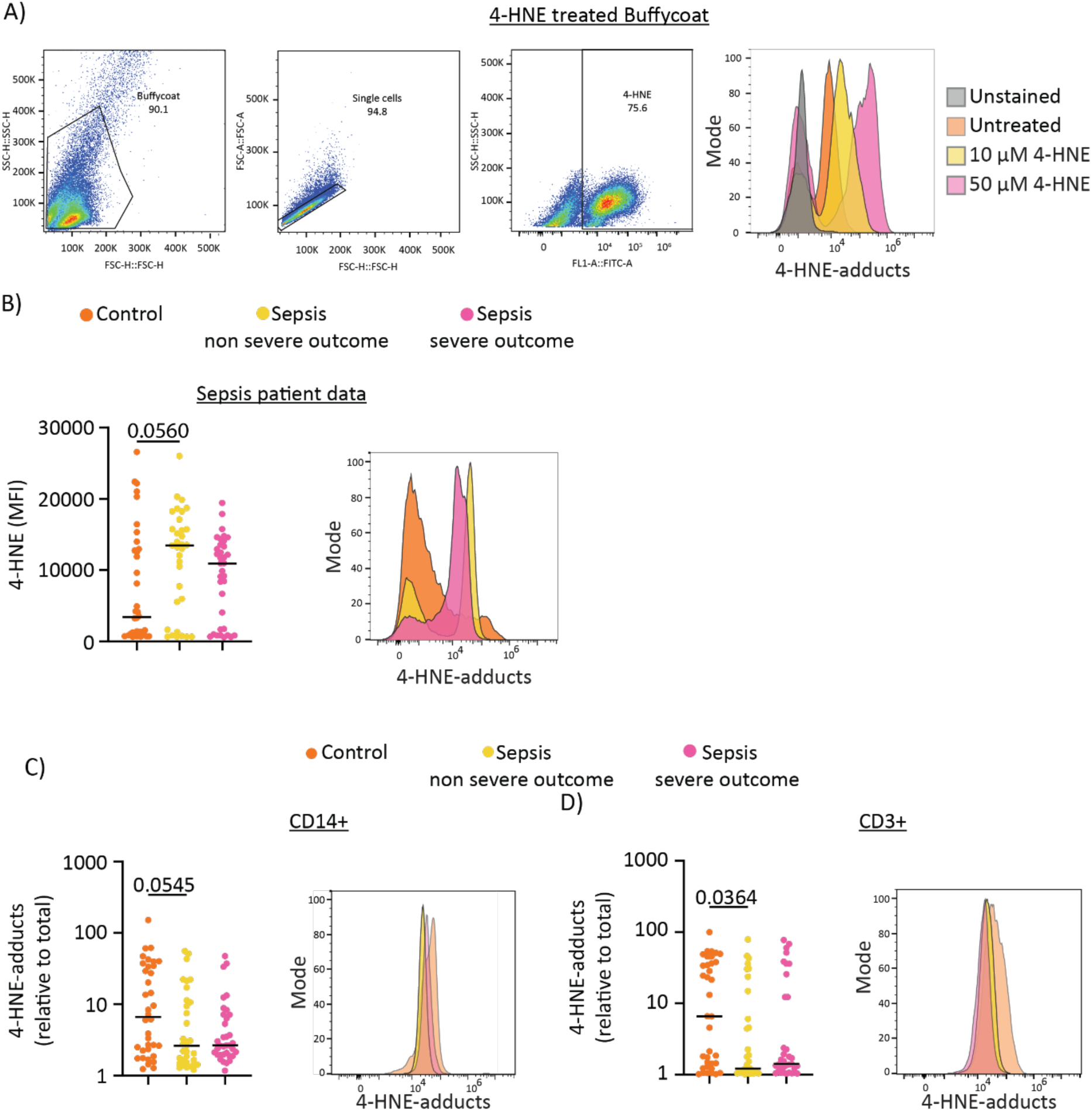
4-HNE protein adducts are increased in Patients with sepsis. Buffy coats were stained with an antibody against 4-HNE protein adducts and measured by flow cytometry. A) Gating strategy of buffy coat from a healthy individual treated with 10 μM or 50 μM 4-HNE. B) 4-HNE protein adduct levels in patients control (orange), sepsis non-severe (yellow), and severe outcomes (pink). Buffy coats were stained with antibodies against 4-HNE protein adducts, CD3 and CD14, and measured by flow cytometry. C-CD) 4-HNE protein adduct levels in CD14^+^ monocytes (B) and CD3^+^ T cells (C) and representative histograms. Each data point represents one patient, and the bar indicates the median. Data were tested for normal distribution using the Shapiro-Wilk test before being analysed with a suitable ANOVA. Histograms for representative patients of each group are shown.

Our flow cytometry approach allowed for discerning 4-HNE protein adduct levels in different immune cell types. Therefore, we co-stained the cells for CD14 and CD3 to determine 4-HNE protein adduct levels, specifically in CD14^+^ monocytes and CD3^+^ T cells. The gating strategy is shown in Supplementary Figure 1. Surprisingly, the levels of 4-HNE protein adducts in the CD14^+^ and CD3^+^ cells were significantly lower in the sepsis patient groups compared to the healthy donors (Figure 1C-D). Thus, increased 4-HNE levels, as observed in buffy coat data, are caused by the CD14^-^CD3^-^ population, which are likely neutrophils (CD15^+^) and/or natural killer (NK) cells (CD3^-^, CD56^+^).

#### IL-10 levels are increased in sepsis and seem to correlate with mortality

IL-10 is produced in bacterial sepsis and is the highest in septic shock (*36*). The production of IL-10 in septic shock positively correlates with the production of pro-inflammatory cytokines, including TNF-α and IL-6 (*37*). To confirm these findings, cytokine levels were determined in the plasma of the same patients as used for the 4-HNE measurements in Figure 1. In line with the literature (*5*), we found that the levels of IL-10 and IL-6 correlate with sepsis, whereas no increase (and even a non-significant decrease) was seen for TNF-α (Figure 2A).

**Figure 2:**
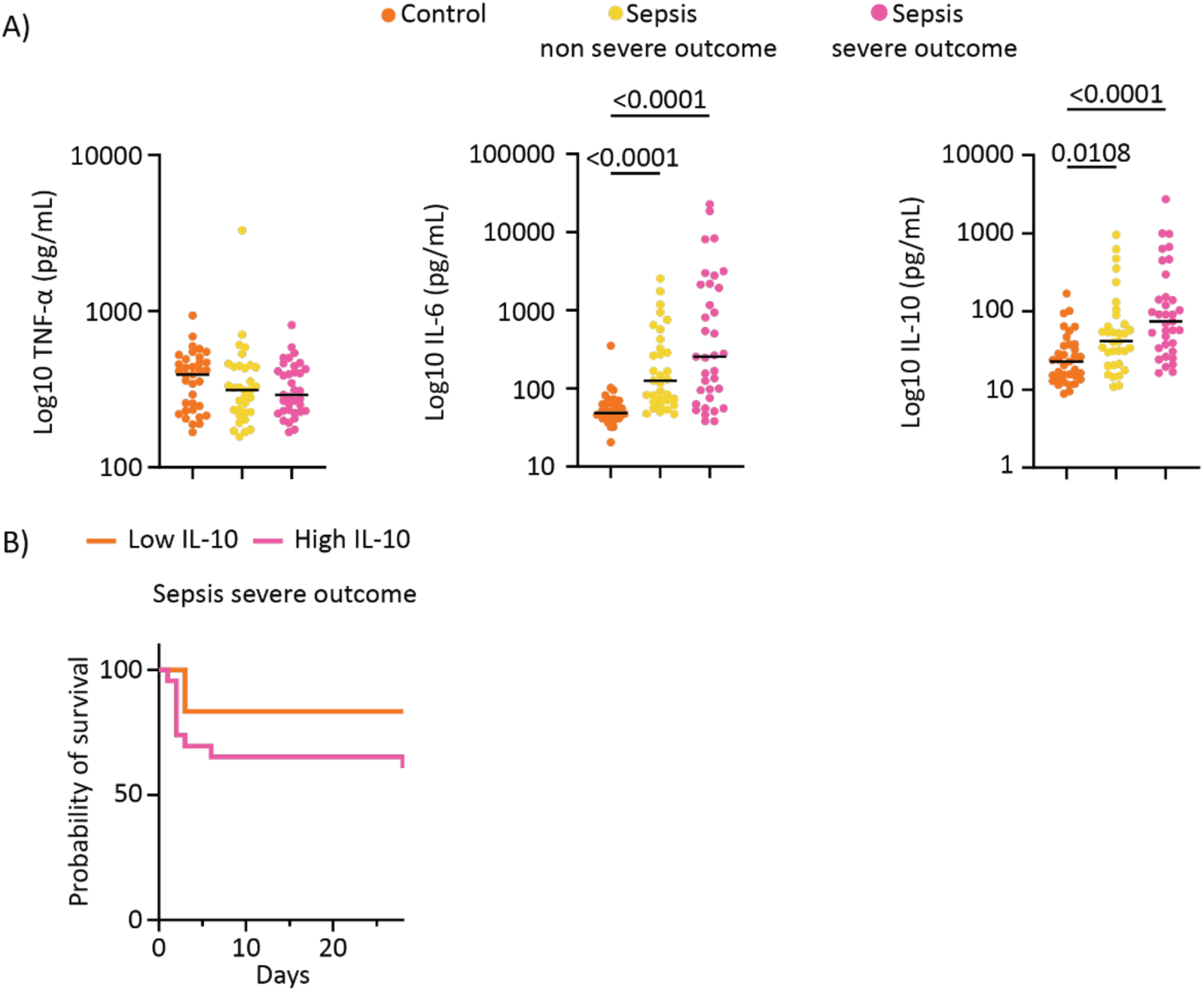
IL-6 and IL-10 are increased in sepsis patients and correlate with sepsis outcome. A) TNF-α, IL-6, and IL-10 levels in the plasma of patients control (orange), sepsis non-severe (yellow), and severe outcomes (pink). Each data point represents one patient, and the bar indicates the median value. Data were analyzed using the Kruskal-Wallis test. B) Kaplan Meier survival curve in different patient groups in case of low IL-10 (<50 pg/mL) and high IL-10 (>50 pg/mL).

High levels of IL-10 production in patients with septic shock correlate with mortality (*6*). To confirm this in our cohort, we plotted the 28-day survival of severe sepsis patients, distinguishing between low and high levels of IL-10 based on the levels observed in non-sepsis patients (<50 pg/mL). Indeed, increased levels of IL-10 correlated with a lower survival rate in the sepsis severe outcome group (Figure 2B), which is in line with previous findings (*6*).

### LPS induces the formation of 4-HNE protein adducts in macrophages

Because 4-HNE affects cellular processes (*21*, *28–30*), we performed experiments with cultured human monocyte-derived macrophages to assess the effect of 4-HNE on cytokine production. We first confirmed that inflammation increases the levels of 4-HNE protein adducts by treating the cells with LPS, which is well known to promote the formation of ROS in human macrophages (*13*). ROS have been reported to increase the formation of 4-HNE in human monocytes and monocyte-derived dendritic cells (*38*). We confirmed that LPS induces ROS production using the fluorescent probe CellROX in human monocyte-derived macrophages treated with 100 ng/mL LPS for 24h (Figure 3A). Albeit significant, the total increase in ROS production was small, possibly because monocyte-derived macrophages have a high baseline of ROS production. To assess 4-HNE protein adduct formation, we also stained LPS-treated macrophages with the antibody recognizing 4-HNE protein adducts. This showed a small but significant increase in the levels of 4-HNE protein adducts in macrophages upon LPS treatment (Figure 3B).

**Figure 3:**
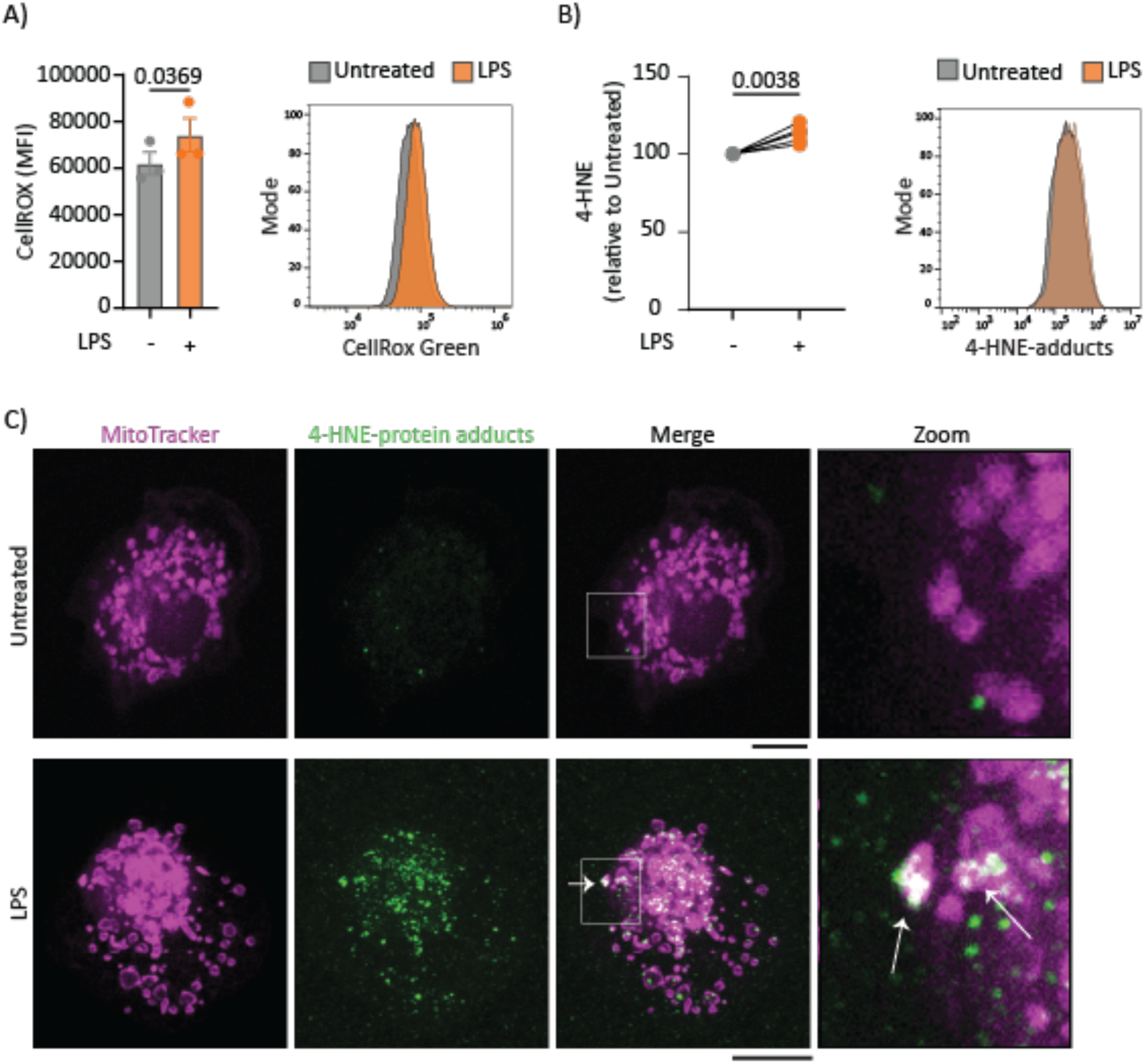
LPS induces 4-HNE production in human monocyte-derived macrophages. Macrophages derived from peripheral blood monocytes of healthy donors were stimulated with 100 ng/mL LPS for 24h. A) Intracellular ROS production was measured with flow cytometry using CellROX Green reagent. Each data point represents one donor (n=3), error bars show SEM, and significance is tested with a paired t-test. The histogram shows untreated (grey) versus LPS (orange) for one donor. B) Macrophages were stained with an antibody against 4-HNE protein adducts and analyzed by flow cytometry. The mean fluorescent intensity is plotted relative to the condition without LPS (grey). Every data point pair represents one donor (n=6), paired t-test. C) Confocal microscopy of untreated and LPS-treated macrophages immunolabeled for 4-HNE protein adducts (green) and co-stained with MitoTracker CMXROS red (magenta). The white arrows show co-localization. Scale bars, 10 μm.

To determine the localization of 4-HNE adducts in macrophages, fluorescence microscopy with immunolabeling was conducted. Co-labeling of the 4-HNE protein adducts with the mitochondrial probe MitoTracker showed that most 4-HNE protein adducts are located in mitochondria (Figure 3C), which is in line with previous findings in adult mouse cardiomyocytes treated with the monoamine oxidase-A (MAO-A) substrate Tyramine (*39*). Mitochondria are an important source of ROS (*40*) and have membranes that contain high levels of carnitine and other ROS-sensitive unsaturated phospholipids, likely explaining the high local 4-HNE levels. Moreover, the microscopy images support the flow cytometry data and showed an increase in 4-HNE protein adduct levels in the LPS-stimulated macrophages compared to untreated cells. Together, these results confirm that LPS-induced ROS provoke an increase in 4-HNE protein adducts, showing that these adducts mainly localize in mitochondria.

#### 4-HNE induces metabolic reprogramming in macrophages

Since LPS treatment results in the accumulation of 4-HNE in mitochondria, the impact of 4-HNE on mitochondrial activity was tested. Human monocyte-derived macrophages were pre-treated with 50 μM 4-HNE for 10 min before being treated for 24h with 100 ng/mL LPS. In this study, all experiments were conducted using complete cell culture media supplemented with 10% fetal bovine serum (FBS), which is more physiological and prevents starvation effects. However, proteins present in FBS can react with 4-HNE (Alviz-Amador et al., 2019; Szapacs et al., 2006), and the effective concentration of non-conjugated 4-HNE that the cells were exposed to can be expected to be much lower than 50 μM. At the 4-HNE concentrations and incubation times used in our study, 4-HNE did not affect the viability of the macrophages (Supplementary Figure 2).

Reorganization of the tricarboxylic acid (TCA) cycle is an important metabolic adjustment that facilitates rapid ATP production and occurs during the activation of macrophages (*41*). LPS provokes an upregulation of aerobic glycolysis, a reduction in metabolic respiration, disruption of the TCA cycle, and an accumulation of succinate (*41*). Our results show the accumulation of 4-HNE in the mitochondria, and 4-HNE has been shown to play a role in inhibiting oxidative phosphorylation (*42*). Therefore, the effect of 4-HNE on mitochondrial activity in macrophages was tested. Our results show a 4-HNE dose-dependent decrease in succinate dehydrogenase (SDH) activity (Complex II) as determined by the MTT (3-(4, 5-dimethylthiazol-2-yl)-2,5-diphenyltetrazolium bromide) assay (Figure 4A).

**Figure 4:**
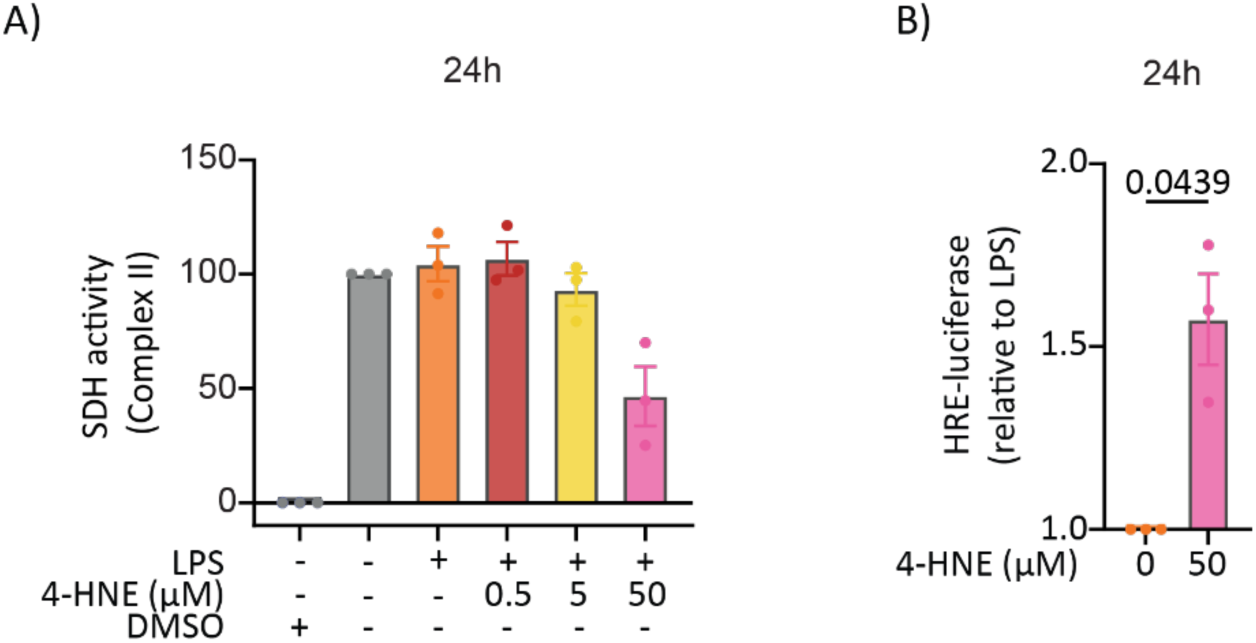
Treatment of macrophages with 4-HNE provokes metabolic reprogramming. Monocyte-derived macrophages were treated with 4-HNE in the presence or absence of LPS. A) Complex II activity was measured after 24h with the MTT assay. Each data point represents one donor, displaying data relative to untreated cells. B) Macrophages were transfected with an HRE-luciferase plasmid to check for HIF-1α activity. Macrophages were stimulated with LPS with or without 4-HNE. Each data point represents one donor, and data are displayed relative to LPS. Error bars display SEM, data analyzed with a paired t-test, and the p-value is indicated.

A reduction in Complex II activity disrupts the electron transfer chain, which can result in the accumulation of succinate due to impaired oxidation to fumarate. Succinate accumulation stimulates activity of the transcription factor HIF-1α and increases mitochondrial ROS production (*43*). To assess the activity of HIF-1α, a hypoxia-responsive element (HRE)-luciferase plasmid was used. Our transfected macrophages were treated with LPS in the presence and absence of 4-HNE. Indeed, the addition of 4-HNE increases HIF-1α activity compared to LPS-stimulated macrophages (Figure 4B).

#### 4-HNE treatment provokes a transcriptional change in macrophages

Our data show that 4-HNE upregulates the activity of the transcription factor HIF-1α, so we performed transcriptional profiling of human monocyte-derived macrophages derived from 8 healthy donors. The cells were pre-treated with 50 μM 4-HNE for 10 min before being treated for 24h with 100 ng/mL LPS. RNA sequencing revealed that the treatment with 50 μM 4-HNE resulted in significant transcriptional reprogramming (Figure 5A). Pathway analysis demonstrated that 4-HNE treatment influences gene expression of multiple signaling pathways, including the HIF-1, MAPK, and NF-κB signaling pathways (Figure 5B). Treatment with 4-HNE provoked the upregulation of 1,178 genes and the downregulation of 1,404 genes compared to macrophages exposed to LPS alone (Figure 5C). When examining the expression levels of cytokine coding and related genes, we found that 4-HNE significantly decreased *IL1RN, IL10*, *PTGS2*, and *NFKBIA* (Figure 5D, Supplementary Figure 3). This effect was absent for other inflammation-related genes, including the genes coding for the inflammatory cytokines IL6 and TNF-α (Figure 5D).

**Figure 5:**
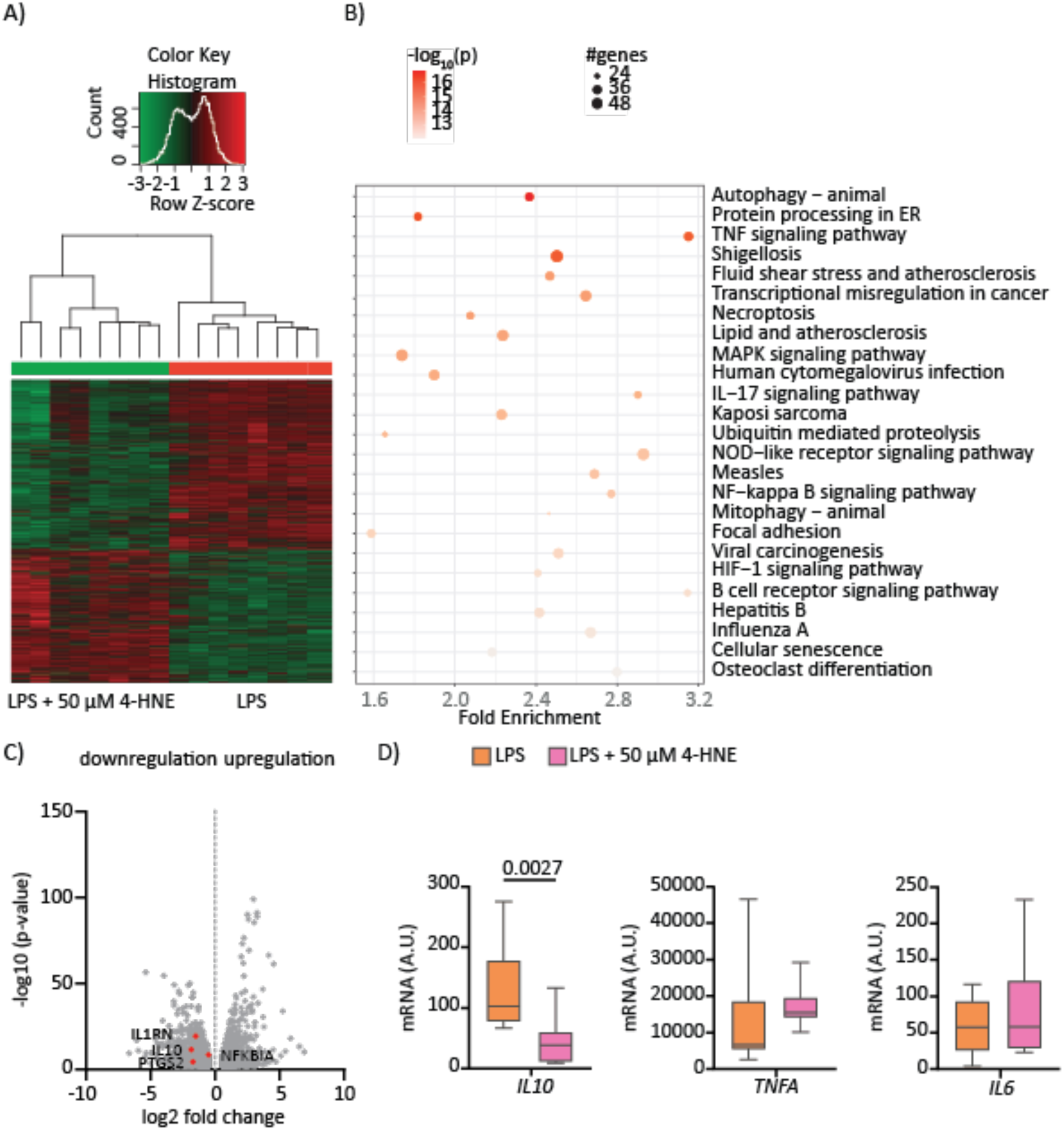
Treatment of macrophages with 4-HNE triggers transcriptional changes. Transcriptomics analysis of human monocyte-derived macrophages from 8 healthy donors treated with LPS or LPS + 50 μM 4-HNE for 24h. A) The heat map displays differential gene expression in LPS (red bar, right panel) vs. LPS+4HNE-treated macrophages (green bar, left panel). B) Pathway analysis shows significant (p<0.05) gene changes in distinct pathways. The size of the circle displays the number of genes that are influenced, whereas the color indicates the -log10 p-value. The x-as shows the fold enrichment, and the y-as shows transcriptionally influenced pathways. C) Vulcano plot highlights genes that are significantly upregulated (right) or downregulated (left) through the treatment with 4-HNE. The red dots highlight distinct genes that are indicated by text. D) RNA sequencing results for specific genes for 8 donors that were treated with LPS (orange) or LPS + 50 μM 4HNE (pink). Data are displayed as arbitrary units of mRNA counts, and significance is tested with a paired t-test.

#### 4-HNE suppresses IL-10 production in macrophages but does not affect IL-6 and TNF-α

The expression levels of several inflammation-related genes were tested by RT-qPCR to validate the results obtained from the RNA sequencing. In accordance with the RNA sequencing results, stimulation with 50 μM 4-HNE and LPS decreased the mRNA levels of *IL10*, *PTGS2*, and *IL1RN* compared to LPS alone, whereas no differences were observed for *IL6* and *TNF* (Figure 6A; Supplementary Figures 3 & 4). Thus, 4-HNE suppresses the transcription of *IL10* and certain other inflammation-related genes in human monocyte-derived macrophages.

**Figure 6:**
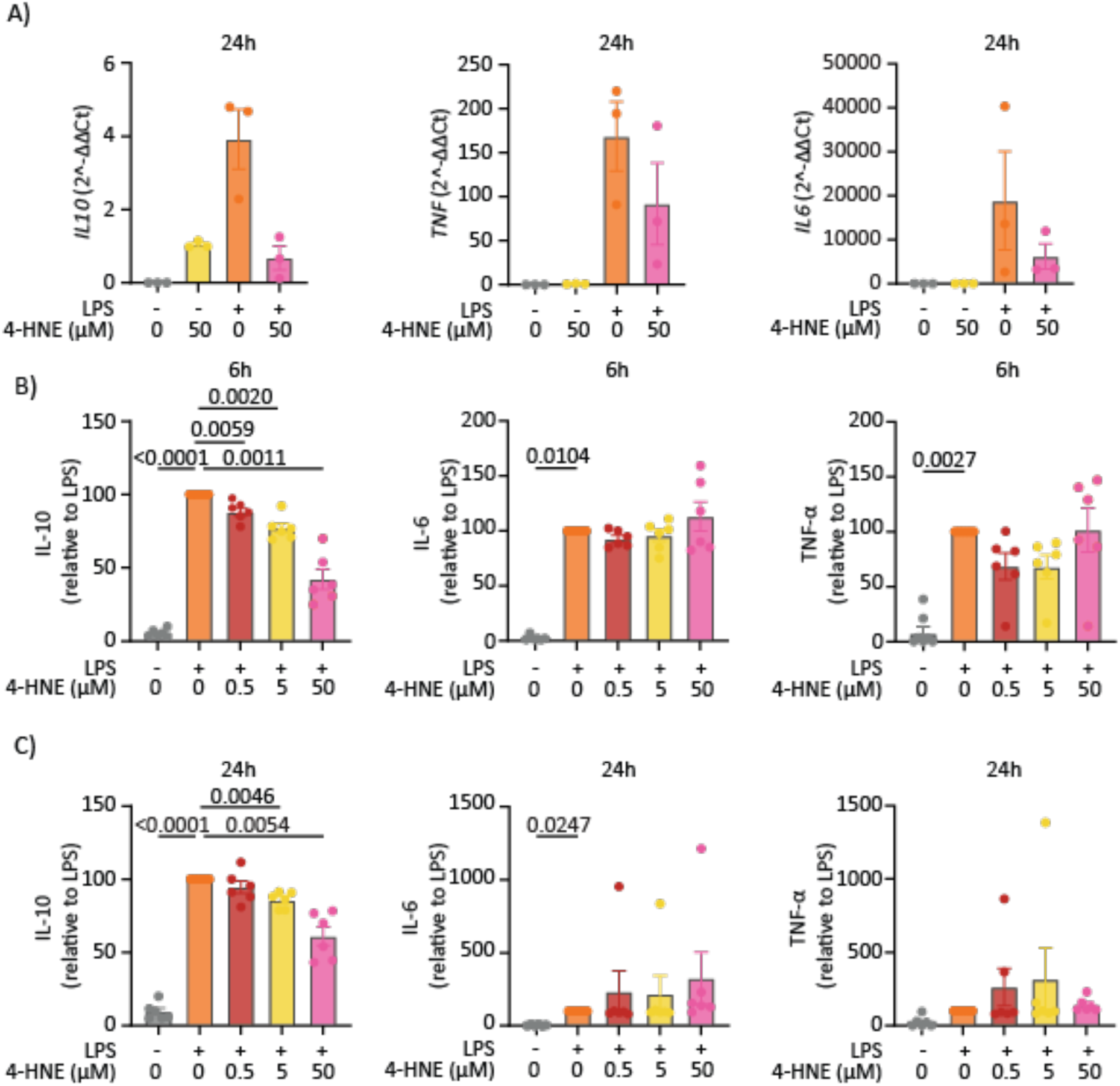
Treatment with 4-HNE suppresses IL-10 production. A) mRNA levels of the indicated genes in monocyte-derived macrophages determined by RT-qPCR. Each data point represents one donor, and the error bars display the SEM. Significance was tested using the RM one-way ANOVA. p-values are displayed in case of significance. Cytokine production from monocyte-derived macrophages measured 6h B) and 24h C) after 4-HNE treatment in the presence or absence of LPS. Each data point represents one donor, and the data are displayed relative to LPS without 4-HNE. Data are analyzed with the RM one-way ANOVA or Friedman test depending on the data distribution, n=6; error bars show SEM.

Next, the effect of 4-HNE on cytokine production at the protein level was determined at 6h and 24h after incubation. In line with our transcriptomics and RT-qPCR data, treatment with 4-HNE resulted in a dose-dependent decrease of IL-10 production after 6h and 24h incubation (Figure 6B-C). Also in line with our RNA sequencing data, no effect of 4-HNE treatment was found for TNF-α and IL-6 (Figure 6B-C), although there was considerable variability in cytokine levels among donors.

#### 4-HNE inhibits STAT3 signaling

Interactions of IL-10 with the IL-10 receptor induce the activation of STAT3 by phosphorylation of tyrosine residue 705. STAT3 is a transcription factor that promotes the expression of anti-inflammatory molecules, such as SOCS3 (*44*). Our RNA sequencing data showed that expression levels of *STAT3* were also reduced by 4-HNE treatment (Figure 7A). In addition, expression levels of *SOCS3* were also decreased upon 4-HNE treatment (Figure 7A).

**Figure 7:**
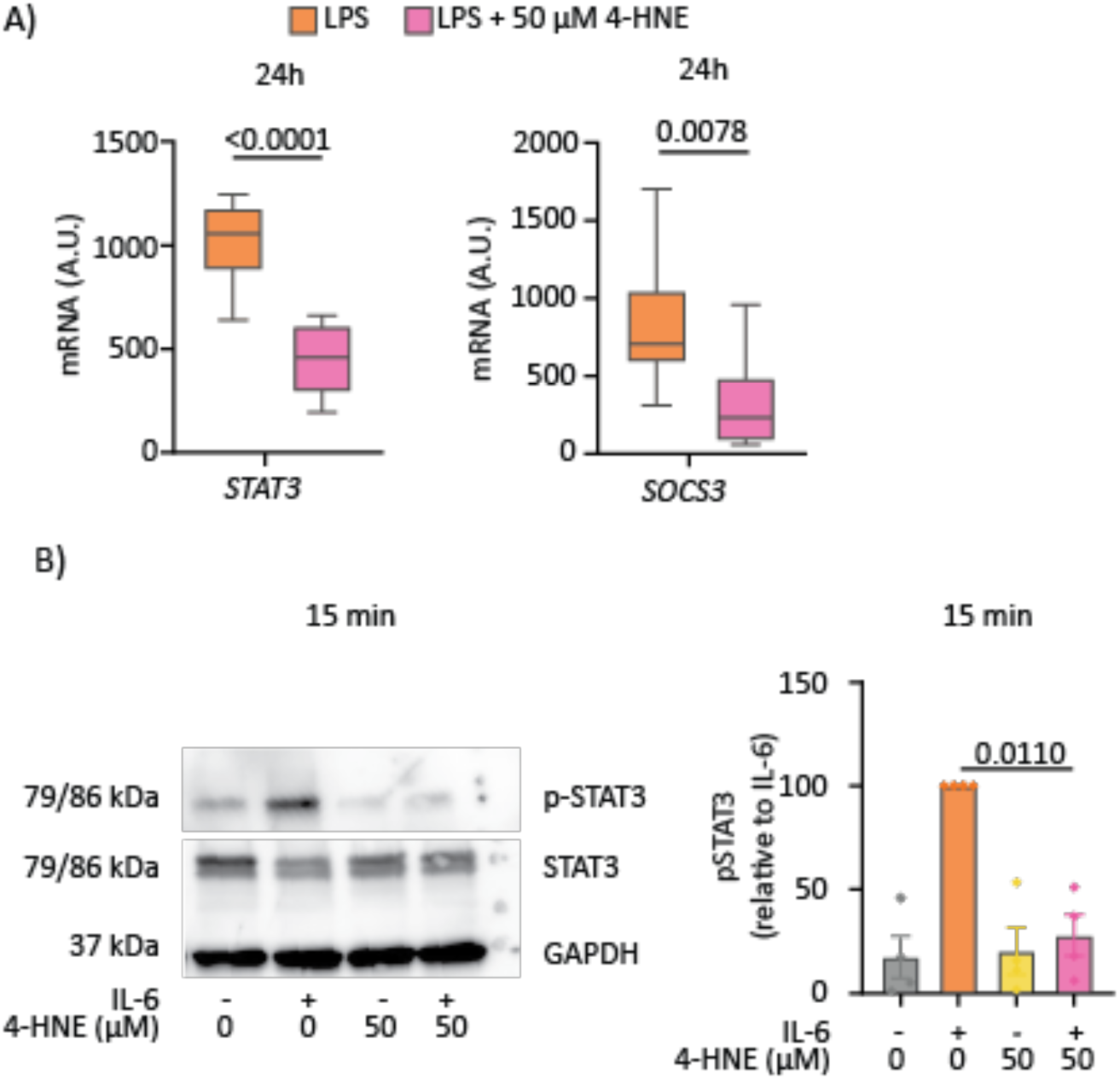
Treatment with 4-HNE suppresses STAT3 signaling. A) RNA sequencing results for *STAT3* and *SOCS3* for macrophages that were treated with LPS (orange) or LPS + 50 μM 4-HNE (pink), n=8, paired t-test. B) Western blot (left) and quantification (right) for pSTAT3, STAT3, and GAPDH protein expression. Data are analyzed with GAPDH as a reference and displayed relative to IL-6. Each data point shows one donor, and data are analyzed with an RM one-way ANOVA, n=3, error bars show SEM.

To further assess the effects of 4-HNE on STAT3 signaling, we treated the macrophages with IL-6 in the presence and absence of 50 μM 4-HNE. For these experiments, we used IL-6 to activate STAT3 (*44*, *45*) instead of IL-10, as this allowed us to discern the effects on STAT3 signaling independently of IL-10. We did not observe a significant effect on total levels of STAT3, however, treatment with 50 μM 4-HNE decreased tyrosine 705 phosphorylation of STAT3 (Figure 7B; Supplementary Figure 5). These results show that 4-HNE suppresses the STAT3 pathway in monocyte-derived macrophages.

#### 4-HNE suppression of IL-10 production can be reversed by glutathione

4-HNE has an estimated half-life of less than 2 minutes due to detoxification mechanisms that neutralize 4-HNE (*46*). One significant detoxification mechanism is the conjugation of 4-HNE with glutathione (GSH), to form GSH-4-HNE conjugates, a reaction that is catalyzed by the enzyme glutathione-S-transferase but can also occur spontaneously (*27*, *46*). To confirm that the reduction in IL-10 production was induced by reactive 4-HNE, human monocyte-derived macrophages were pre-treated for 1h with different concentrations of GSH before adding 4-HNE and LPS. TNF-α, IL-6, and IL-10 production were measured after 6h and 24h treatment (Figure 8). As we observed previously, treatment with 50 μM 4-HNE and LPS reduced the IL-10 production by approximately 70% compared to LPS alone. However, this effect could be blocked in a dose-dependent fashion by GSH. In contrast, detoxification of 4-HNE by GSH did not affect the production of TNF-α and IL-6. These results confirm that 4-HNE needs to be reactive to block IL-10 production, as the detoxified conjugate did not affect LPS-induced IL-10 production.

**Figure 8:**
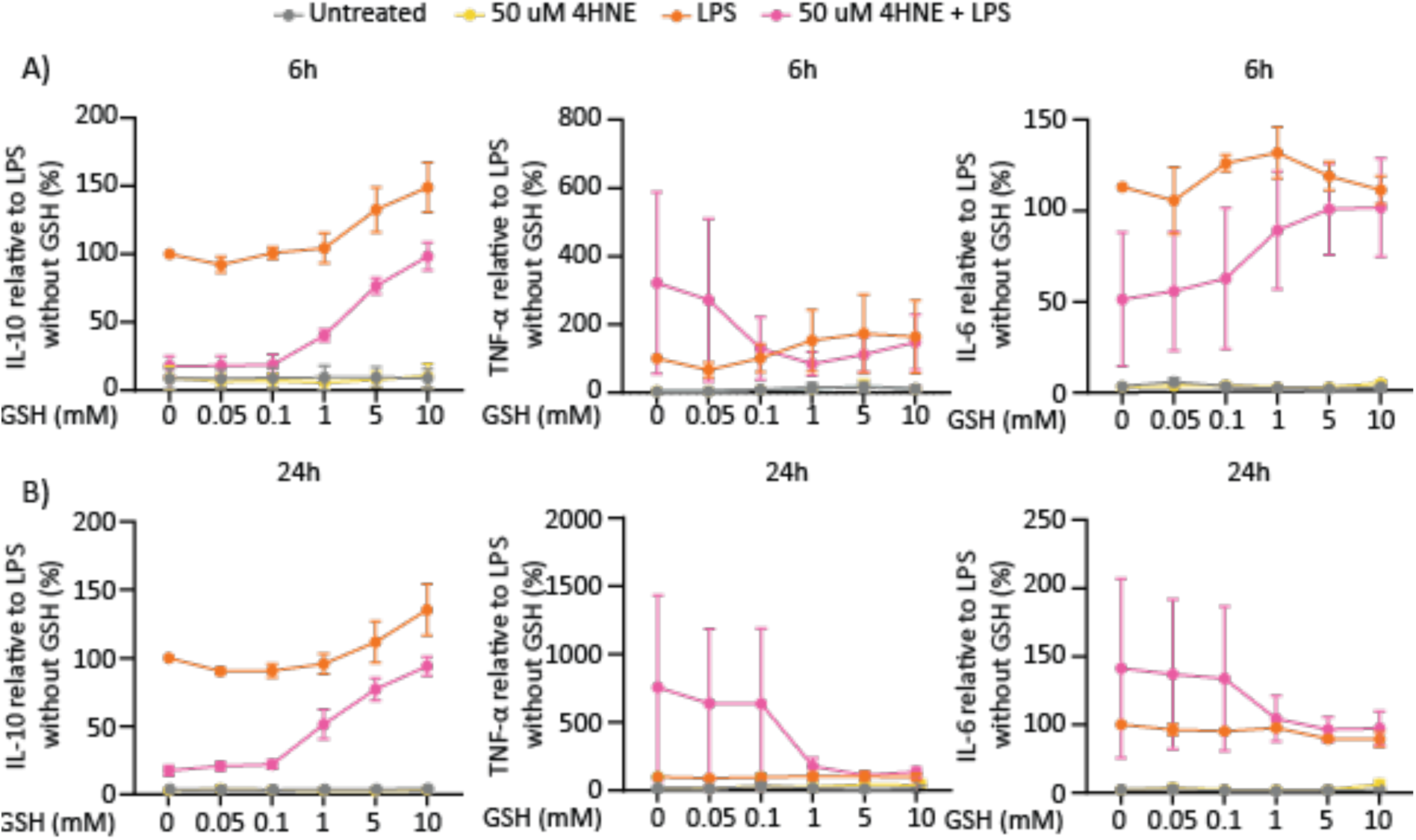
Glutathione blocks 4-HNE-mediated inhibition of IL-10 production. Cells were pre-treated with the indicated concentrations of glutathione prior to 4-HNE and LPS stimulation. Cytokine production was determined by ELISA after 6h (A) and 24h (B). Each data point shows the average from three donors. The error bars indicate the SEM. Data are plotted as a percentage compared to LPS without glutathione.

#### 4-HNE treatment decreases NF-κB and stimulates p38 activity

LPS stimulation of macrophages triggers multiple signaling pathways that promote the production of TNF-α, IL-6, and IL-10, including the main inflammatory pathways p38 and NF-κB. We therefore assessed the effects of 4-HNE on these pathways to explore how it selectively blocks LPS-induced IL-10 production. Since our RNA sequencing indicated that 4-HNE affects NF-κB signaling (Figure 5B), we first assessed this pathway. We used a human monocyte THP-1 reporter cell line containing a stable NF-κB inducible secreted alkaline phosphatase (SEAP) reporter construct. Upon activation of NF-κB, this cell line produces SEAP, which can be measured in the supernatant using a chromogenic substrate conversion assay (*47*). Using the same experimental setup as in our previous experiments, the cells were treated with different concentrations of 4-HNE followed by LPS for 6h, 24h, and 48h (Figure 9A). 50 μM of 4-HNE reduced NF-κB activity after 6h and 24h incubation, in line with previous results in human monocytes (*48*). The effect was no longer observed after 48h, possibly because all 4-HNE was reacted away. Thus, 4-HNE reduces LPS-induced NF-κB signaling. This is striking, as we did not observe any effects of IL-6 or TNF-α in our cell models. Therefore, we investigated the possibility of a compensatory mechanism that drives the secretion of IL-6 and TNF-α.

**Figure 9:**
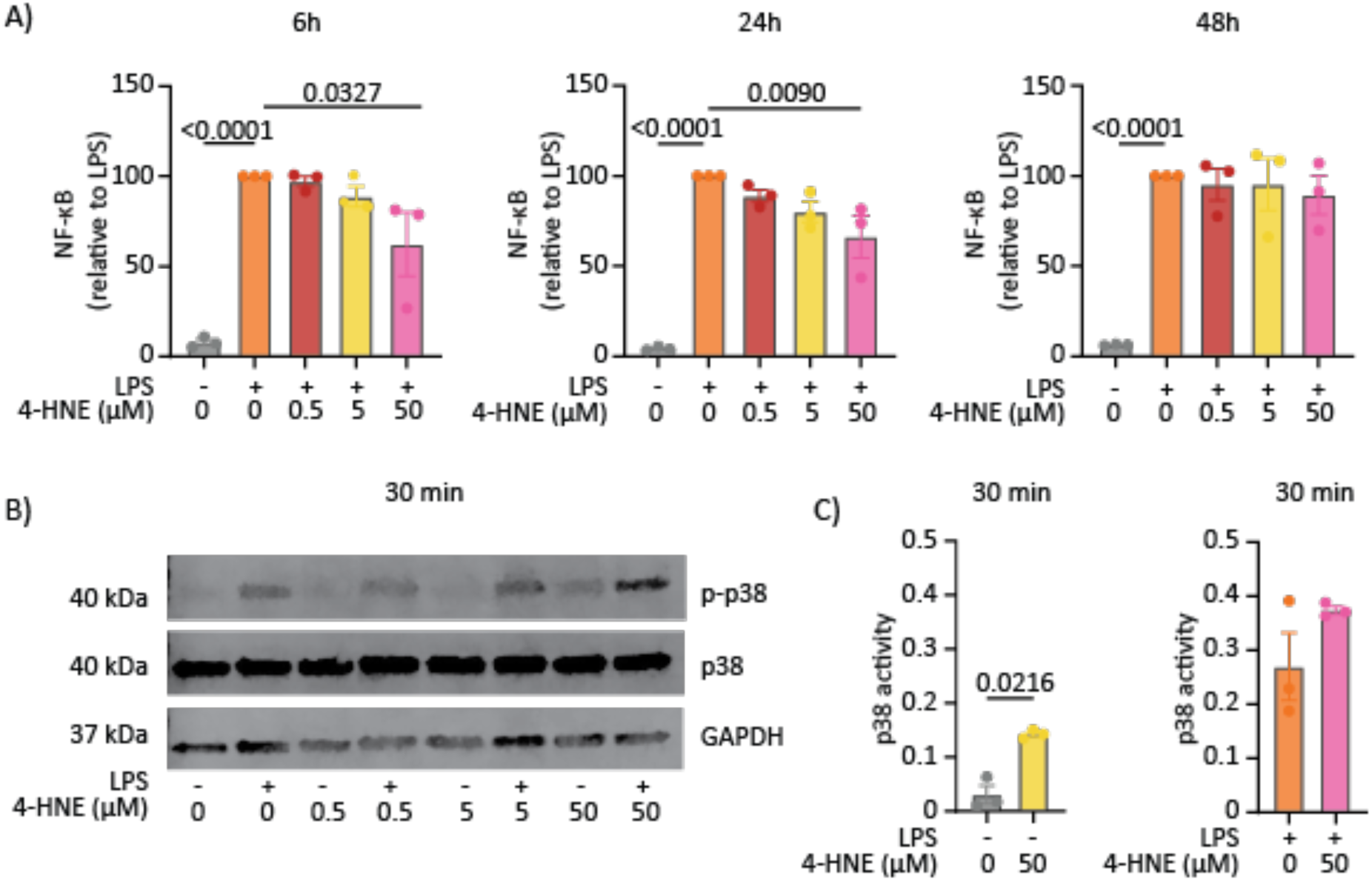
4-HNE inhibits NF-κB and promotes p38 signaling. A) The human monocyte NF-κB reporter cell line Thp1 was treated with different concentrations of 4-HNE in the presence or absence of LPS. NF-κB activity was determined after 6h, 24h and 48h. The experiment was done in triplicates and repeated three times. Data were analyzed by Ordinary one-way ANOVA, error bars show SEM. B-C) P38 activity was determined by western blot of phosphorylated p38 (40 kDa), total p38 (40 kDa), and GAPDH (37 kDa, loading control). Representative blot (B) and quantification of phosphorylated over total p38 (C) are shown. Each data point represents one donor, n=3, paired t-test.

Our RNA sequencing also revealed that MAPK signaling was affected by 4-HNE (Figure 5B). Effects of 4-HNE on p38 MAPK have been reported previously. For example, in the murine macrophage cell line J774A.1 treatment with 10 μM 4-HNE increased p38 activity, with the highest activation after 30 and 45 min of treatment (*49*). Another study showed that stimulation of the human monocyte cell line THP-1 with 20 μM 4-HNE also induced p38 activation, which was reduced by adding the antioxidant N-acetylcysteine (*50*). To confirm these findings in our cell type, the effect of 4-HNE on p38 phosphorylation at activation sites Thr180/Tyr182 was investigated in human monocyte-derived macrophages treated with 4-HNE in the presence or absence of LPS for 30 min. 50 μM of 4-HNE increased phosphorylation of p38 in the absence and presence of LPS (Figure 9B-C; Supplementary Figure 6). This effect did not reach significance in the presence of LPS due to donor variation.

#### 4-HNE or p38 inhibition alone suppress IL-10, whereas IL-6 and TNF-α suppression require both

Given our findings that 4-HNE selectively suppresses IL-10 but did not affect IL-6 and TNF-α production and that 4-HNE oppositely affects NF-κB (inhibition) and p38 MAPK (increase), we investigated whether the differential effects of 4-HNE on cytokine production could be due to a different reliance on these signaling pathways. Thus, we hypothesized that IL-10 production would require both NF-κB and p38 signaling, whereas for IL-6 and TNF-α production, increased p38 signaling could compensate for a reduction of NF-κB signaling. To test this, macrophages were pre-treated with the p38 inhibitor SB302580, and cytokine production was tested after 6h and 24h incubation with LPS and/or 4-HNE.

As predicted by our hypothesis, SB302580 decreased IL-10 production in LPS-treated cells, comparable with the inhibition with 4-HNE (Figure 10A). This reduction of IL-10 production was stable up to 24h. However, also in accordance with our hypothesis, the production of TNF-α and IL-6 were not affected by SB302580 or 4-HNE treatment alone, whereas they were decreased following treatment with a combination of both compounds (Figure 10B-C). These data thus show that IL-10 production requires both p38 and NF-kB signaling, whereas for IL-6 and TNF-α production p38 signaling can compensate for a reduced NF-kB signaling. In addition, the combined blockage of p38 with SB302580 and NF-kB with 4-HNE is required to inhibit IL-6 and TNF-α production.

**Figure 10:**
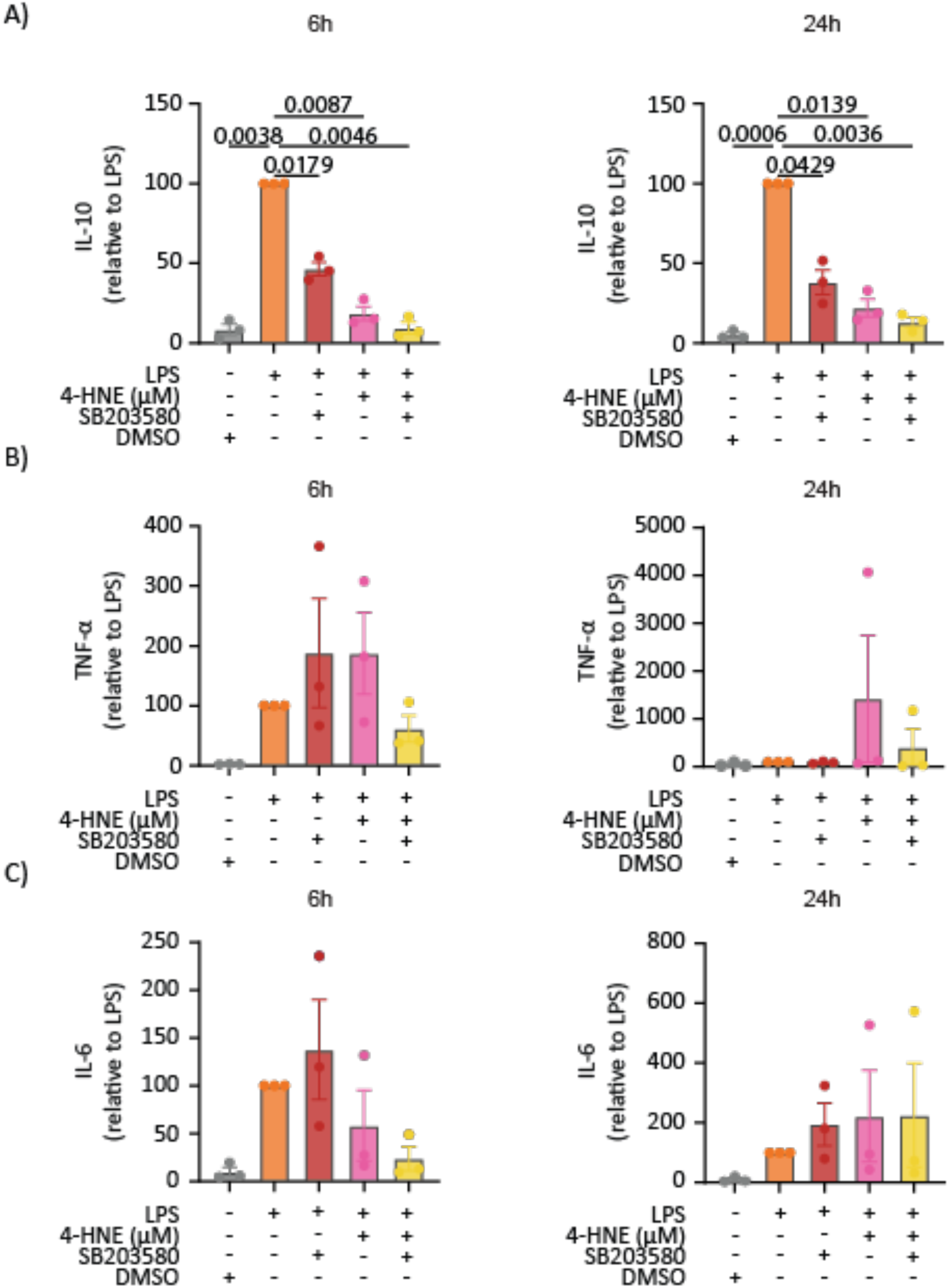
4-HNE or p38 inhibition alone reduce IL-10, whereas IL-6 and TNF-α reduction require both. Monocyte-derived macrophages were pre-treated with p38 inhibitor SB320580 prior to treatment with 4-HNE and LPS. A-C) IL-10 (A), TNF-α (B) and IL-6 (C) levels were determined by ELISA at 6h (left) or 24h (right) after LPS. Data are displayed relative to LPS, and each data point represents one donor. The error bar shows the SEM. Data are analyzed using Friedman or an RM one-way ANOVA test, depending on the distribution of the data points.

## Discussion

ROS are elevated in sepsis and correlate with severe outcome (*35*, *44*, *51*). Elevated ROS promote the production of reactive aldehydes, including 4-HNE. As 4-HNE can dose and cell type dependently influence the immune response (*52–57*), we determined its link to sepsis.

In our study, we measured the levels of 4-HNE protein adducts in the buffy coats of patients with sepsis, with and without severe outcomes, and acutely ill patients without an infection served as the control group. The levels of 4-HNE protein adducts were lower in the control group compared to the sepsis groups. This finding aligns with the previously reported increased levels of ROS and MDA in sepsis (*35*). In accordance with this, the stimulation of cultured macrophages with LPS increased both ROS and 4-HNE protein adduct levels, as also observed previously (*13*, *58*). Our immunostaining revealed that most 4-HNE adducts are present in mitochondria, likely because mitochondria are the main sites of ROS production (*59*) and contain high levels of PUFAs that are prone to oxidation by ROS (*60*). This is potentially relevant for sepsis, as impaired mitochondrial dysfunction and uncoupling of mitochondrial respiration are hallmarks of sepsis (*31*, *32*). Systemic inflammation likely demands high energy, driving hypermetabolic mitochondria to increase energy production and support processes such as fever, elevated protein synthesis, and increased heart and respiratory rates. Consequently, patients with sepsis may require more energy, which results in higher metabolic activity to generate more ATP, resulting in an increase in ROS generation (*61*). Since ROS induce cell damage leading to an increase in lipid peroxidation (*35*), this likely explains the accumulation of 4-HNE protein adducts in mitochondria.

Metabolic reprogramming influences the immune response in macrophages. Potentially, 4-HNE contributes to immune dysregulation in sepsis patients through IL-10 due to metabolic reprogramming. A previous study has shown that a decrease in complex II (SDH) activity leads to an increase in ROS production by reverse electron transport from complex I and an accumulation of succinate, a metabolite that regulates major inflammatory pathways (*43*). An increase in succinate prevents the degradation of HIF-1α by competitive binding to the prolyl hydroxylase domain (PHD) (*62*). In LPS-stimulated bone-marrow-derived macrophages, increased succinate stimulated HIF-1α activity, causing increased IL-1β and decreased IL-10 production, but did not affect TNF-α (*43*). This aligns with our results, where supplementation of LPS-stimulated macrophages with 4-HNE led to a decrease in complex II activity and an increase in HIF-1α activity paired with a strong and dose-dependent reduction in IL-10 production and downstream STAT3 activation. In contrast, TNF-α and IL-6 production were unaffected. These effects could be blocked with glutathione, an antioxidant that reacts with 4-HNE (*27*, *46*), showing that the reactivity of 4-HNE is important for these effects.

Mechanistically, transcriptomics analysis showed that 4-HNE in LPS-treated macrophages altered the expression of many genes involved in inflammatory signaling pathways regulating IL-10 production, including p38 MAPK and NF-κB. 4-HNE oppositely affects these signaling pathways, as we found that it stimulates MAPK p38 activation but reduces NF-κB activation. Although we could not reproduce previous findings in human peripheral blood monocytes, where 4-HNE inhibits TNF-α production (*48*), potentially due to differences in handling procedures, our findings align with findings in other studies. In the murine macrophage cell lines RAW264.7 and J774A.1, treatment with 4-HNE promotes p38 activation (*29*, *49*). Moreover, 4-HNE reduces NF-kB signaling in primary rat Kupffer cells (*63*), human monocytes (*48*), rat vascular smooth muscle cells, and HepG2 cells (*64*, *65*). However, the inhibition of NF-κB seems cell type, time, and/or concentration specific, as 4-HNE has also been reported to stimulate NF-kB signaling in human fibroblasts, RAW264.7 cells, and smooth muscle cells (*66–69*).

As our data show that 4-HNE or p38 inhibition alone suffices to suppress IL-10 production, whereas only the combination of 4-HNE and p38 inhibition blocks IL-6 and TNF-α production, we propose that the inverse regulation of these pathways could explain the observed specific blockage of IL-10 production. Treatment of macrophages with 4-HNE blocks NF-κB and increases p38 signaling, which results in less IL-10 expression, whereas for IL-6 and TNF-α, this effect might be compensated for by the increase in p38 activation. As we previously showed, HIF-1α can stimulate p38/MAPK through an upregulation of MAP3K8 in monocyte-derived dendritic cells (*70*) and HIF-1α suppressed NF-κB signaling in HeLa cells and *Drosophila melanogaster* (*71*). It should be noted that our experiments were performed in the presence of 10% FBS, which contains many proteins like albumin, globulins, and growth factors that can interact with 4-HNE (*72*). Therefore, the effective concentration of reactive 4-HNE that our cells were exposed to can be expected to be significantly lower due to sequestration by proteins in the FBS.

As an increase in lipid peroxidation (*35*) and mitochondrial dysfunction (*31*, *32*) are hallmarks of sepsis, we investigated the role of 4-HNE in patients’ blood samples. Not only did we find that sepsis patients had the highest plasma levels of IL-10, a finding consistent with literature (*36*, *73*), but importantly, even though total 4-HNE protein adduct levels measured in buffy coat were higher in sepsis patients, they were lower in two main IL-10-producing cell types: CD14^+^ monocytes and CD3^+^ T cells (*74*). The lower levels of 4-HNE in monocytes and T cells likely facilitate these cells’ elevated production of IL-10. The reason for this cell-type-specific reduction is unclear, but given that we find that most 4-HNE is located in mitochondria, it might be that the mitochondria are altered in monocytes and T cells of sepsis patients. Although the reasons for this are unclear, one possibility is that T cells and monocytes can produce less ROS (and hence less 4-HNE) because of a lack of oxygen. Neutrophils are far more abundant in blood than monocytes and T cells, and neutrophils also produce higher levels of ROS than other immune cell types; we previously showed that neutrophils produce 10-fold more ROS than monocyte-derived macrophages upon LPS stimulation (*13*). As a consequence, the massive conversion of oxygen into ROS by neutrophils might limit the oxygen availability for ROS (and 4-HNE) production by the T cells and the monocytes, although this remains an untested hypothesis.

IL-10 has been considered a diagnostic target for sepsis (*75*), as the levels of IL-10 in septic shock correlate positively with the intensity of the inflammatory response (*76*). Moreover, IL-10 is also a target for therapeutic intervention because it limits the host’s immune response to pathogens and could prevent organ damage (*77*). Indeed, experiments in IL-10 knockout mice showed that they are more susceptible to CLP-induced sepsis mortality, and this could be rescued by administering recombinant human IL-10 (*78*). Moreover, studies in a CLP-induced sepsis mouse model found that intrathymic injection of a recombinant adenovirus expressing human IL-10 could improve survival rates (*79*). This study showed that only local elevation of IL-10 in the thymus provided protection against mortality because intravenous administration had no impact on outcomes despite preventing the systemic pro-inflammatory response (*79*). In IL-10-deficient mice, the local thymic expression of human IL-10 also improved survival in CLP-induced sepsis (*80*). These results show the beneficial effect of IL-10, whereas we and others (*6*) show that high IL-10 is correlated with mortality. One explanation for this discrepancy could be that patients with sepsis who produce high levels of pro-inflammatory cytokines, such as IL-6, also release more IL-10 in response to systemic inflammation. In this case, the IL-10 does not itself cause the high mortality but is a consequence of high systemic inflammation.

Next to IL-10, our data suggest that 4-HNE could serve as an amenable biomarker for precision medicine in sepsis. Our data show that whereas total levels of 4-HNE protein adducts are increased in sepsis, the levels on monocytes and T cells are lower and inversely correlate with IL-10 production. 4-HNE is strongly immunomodulatory and almost completely blocks IL-10 production *in vitro* but does not affect the production of IL-6 and TNF-α. Therefore, lowering the levels of 4-HNE production might be a new approach to selectively increase IL-10 levels without affecting other inflammatory cytokines. In line with this hypothesis, intravenous injection of an anti-4-HNE antibody in LPS-challenged mice prevented liver injury and death (*81*). Thus, 4-HNE’s immunomodulatory properties, especially its selective inhibition of IL-10 production, make it a potential therapeutic candidate for inflammatory diseases like sepsis.

## Methods

### Clinical study design and obtaining blood samples from patients

Acutelines is a data biobank at the Emergency Department (ED) of the University Medical Center Groningen (UMCG) that prospectively collects data and biomaterials from acutely ill patients. The ED nurse and a researcher screen patients for eligibility upon arrival at the ED. Upon triage, before initiation of treatment, blood samples were collected in EDTA tubes, centrifuged for 15 min at 1,500 x g, and stored at -80°C in the Acutelines biobank. Bed-side data monitored (i.e., vital parameters) are automatically captured and stored, and information from other data sources, such as the electronic health records of the hospital, emergency medical services- and the general practitioner, the municipal registration, health insurance companies, and the pharmacy, is securely imported. De novo research data were collected and managed using REDCap electronic capture tools hosted at the UMCG. All participants gave (deferred) informed consent (by proxy) on enrolment. In case the patient or proxy could not reasonably be reached or when only routine data for clinical care were collected, an opt-out procedure was in place. At www.acutelines.nl the complete protocol and overview of the actual, full data dictionary are available. A detailed description of the design, inclusion, and exclusion criteria has been published previously (*82*). This work has been carried out following The Code of Ethics of the World Medical Association (Declaration of Helsinki) for experiments involving humans. The Medical Ethics Review Committee (METC) of the UMCG reviewed and approved this study (METc; PaNaMa number 18642).

For this study, we post-hoc selected patients who visited the ED for a medical, non-infectious reason (Supplementary table 1), were not admitted to the ICU, and did not die during hospital admission (n=35, control group). Next, we selected patients with sepsis based on a probable or microbiologically confirmed acute bacterial infection from the urinary, respiratory tract, or abdomen as determined by post-hoc adjudication combined with a rise of the SOFA score of two or more within 72 hours as compared to the value measured at the ED. Of these sepsis patients, we selected 35 patients without a severe outcome who were not admitted to the ICU within 72 hours and survived until hospital discharge, and 35 patients with a severe outcome who either were admitted to the ICU or died within 72 hours. All groups were matched based on age and gender.

### Determination of 4-HNE protein adducts and cytokine levels in leukocytes

The buffy coat was used to detect 4-HNE protein adducts in total cells, CD14^+^ monocytes, and CD3^+^ T cells. The buffy coats were thawed and washed several times in cold PBS. The cells were stained with anti-human CD3 antibody (Miltenyi, REAfinity 130-113-141, 1:50) and CD14 antibody (Miltenyi, REAfinity 130-110-583, 1:50) for 10 min at 4°C. Subsequently, cells were washed with cold PBS before fixation with 4% paraformaldehyde (PFA) (Aurion, 15710) for 15 min at 4°C. After fixation, the cells were washed several times in cold PBS, followed by blocking and permeabilization in PBS containing 2% human serum (Sigma/MERCK, H4522-20ML) and 0.05% saponin (Sigma, 47036-50G) for 30 min at 4°C. Subsequently, cells were incubated for 1 hour with primary antibody for 4-HNE protein adducts (Invitrogen, MA5-27570, 1:60) in 2% human serum (Sigma/MERCK, H4522-20ML) and 0.05% saponin at 4°C. Cells were washed three times again with PBS containing 2% human serum and 0.05% saponin before being incubated with secondary antibody donkey-anti mouse IgG (H&L) Alexa flour 488 (ThermoScientific, 10544773, 1:400) for 30 min at 4°C. Finally, the cells were washed twice in PBS before analyzing the samples using a CytoFlex S (Beckman Coulter). Data were analyzed using NovoExpress Software (Agilent).

### Culturing of human monocyte-derived macrophages

Buffy coats from healthy donors were obtained from the Dutch blood bank, who provided informed consent for their use in research. The samples were anonymized, and the investigators could not determine the identity of the blood donors.

PBMCs were extracted from the buffy coats using standard density gradient centrifugation with Lymphoprep (STEMCELL Technologies, 07861) as described earlier (*83*). Monocytes were subsequently separated from the PBMC fraction by magnetic-activated cell sorting (MACS) using CD14 microbeads (Miltenyi Biotec, 130-114-976). The isolated monocytes were cultured in ultra-low adherent 6-well plates (Corning, CLS3471-EA) for seven days with 2 mL of complete media consisting of RPMI 1640 (Gibco, 11530586), 10% fetal bovine serum (FBS) (Hyclone, 10309433), 2 mM L-glutamine (Gibco, 15430614), 1% Antibiotic-Antimycotic (Gibco, 15240062) enriched with 100 ng/mL recombinant human M-CSF (R&D systems, 216-MC) at 37°C and 5% CO_2_. After differentiation, the cells were washed with room temperature PBS and incubated with cold PBS at 4°C for 30 min to detach macrophages. Macrophages were pre-treated with different concentrations of 4-HNE (Sigma/MERCK, 393204) for 10 min at 37°C in complete RPMI. Subsequently, cells were stimulated with 100 ng/mL LPS (Sigma, L4391-1MG) for different time points at 37°C.

### Cell experiment

Macrophages were pre-treated with different concentrations of 4-HNE (Sigma/MERCK, 393204) for 10 min at 37°C in complete RPMI. Subsequently, cells were stimulated with 100 ng/mL LPS (Sigma, L4391-1MG) for different time points at 37°C.

### Confocal microscopy

To determine 4-HNE protein adduct localization, macrophages were seeded on glass coverslips at 50,000 cells/coverslip density. Cells were treated with 200 nM MitoTracker Red CMXRos (Fisher Scientific, 11569106) for 15 min at 37°C and subsequently fixed by incubating cells with 4% PFA (Aurion, 15710) for 15 min at 4°C. Cells were washed three times with PBS and subsequently blocked and permeabilized for 30 min at 4°C using Confocal laser scanning microscopy (CLSM) buffer containing PBS, 20 mM glycine (BoomBv/Acros Organics, 220910010), 3% BSA (Merck, 126575), and 0.1% saponin (Sigma, 47036-50G). Thereafter, cells were stained overnight at 4°C with a primary antibody for 4-HNE protein adducts (Invitrogen, MA5-27570, 1:200) in CLSM with 0.1% saponin (Sigma, 47036-50G) at 4°C. The next day, the cells were washed three times with PBS and 0.1% saponin (Sigma, 47036-50G) and incubated with secondary antibody donkey-anti mouse IgG (H&L) Alexa fluor 488 (ThermoScientific, 10544773, 1:400) for 30 min at room temperature in CLSM with 0.1% saponin (Sigma, 47036-50G). Subsequently, cells were washed three times with PBS and 0.1% saponin (Sigma, 47036-50G), and the coverslips were mounted on glass slides in 70% glycerol containing 1 mM Trolox (Sigma Aldrich, 238813), 100 mM Phosphate buffer (pH 8.0), and 0.33 μg/ml DAPI (Sigma Aldrich, 32670). Samples were imaged using an LSM800 Zeiss confocal laser-scanning microscope with a 63× 1.4 NA oil lens.

### RNA sequencing

Macrophages incubated with and without 4-HNE, followed by LPS stimulation (as described above), were washed three times with cold PBS before lysing them with 1 thioglycerol/homogenization solution. The RNA sequencing facility at UMCG isolated, prepped, and sequenced the 3’end mRNA (5M reads/sample). Next, we performed read mapping and differential expression analysis of the RNA sequencing data. Therefore, RNA-seq data was aligned to the hg38 assembly of the human genome using STAR v2.7.10b (*84*)(parameters: --outFilterMultimapNmax 20 --alignSJoverhangMin 8 --alignSJDBoverhangMin 1 --outFilterMismatchNmax 999 --outFilterMismatchNoverReadLmax 0.04 -- alignIntronMin 20 --alignIntronMax 1000000 --alignMatesGapMax 1000000). Gene counts were computed using featureCounts v2.0.0 (*85*) and the latest refFlat annotation from UCSC (parameters: - M -p). Differential expression analysis was performed using DESeq2 v1.40.1. (*86*). Pathway enrichment analysis was performed using the package pathfindR v2.2.0. (*87*).

### Reverse transcription-quantitative polymerase chain reaction (RT-qPCR)

We performed RT-qPCR to validate the RNA-seq data for selected genes. Therefore, RNA was isolated using a Quick-RNA MiniPrep kit (ZymoResearch, R1054) in accordance with the manufacturer’s instructions. Isolated RNA was used to generate cDNA using the M-MLV-RT kit (Invitrogen, 28025-021) following manufacturer guidelines. cDNA (2 ng/μL per reaction) was used for qPCR using 5 μL SYBR Green Master Mix (Applied Biosystems, A25742) and 2 μL primer mix (FwD + RV) as indicated in Table 3. qPCR was performed using the StepOnePlus Real-Time PCR System (ThermoFisher) using the following protocol: 50°C for 2 min, 95°C for 2 min, 95°C for 15 s, and 60°C for 1 min. This protocol was repeated 40 times. The results were analyzed using the 2-ΔΔCt method as described earlier (*88*).

**Table 3.**
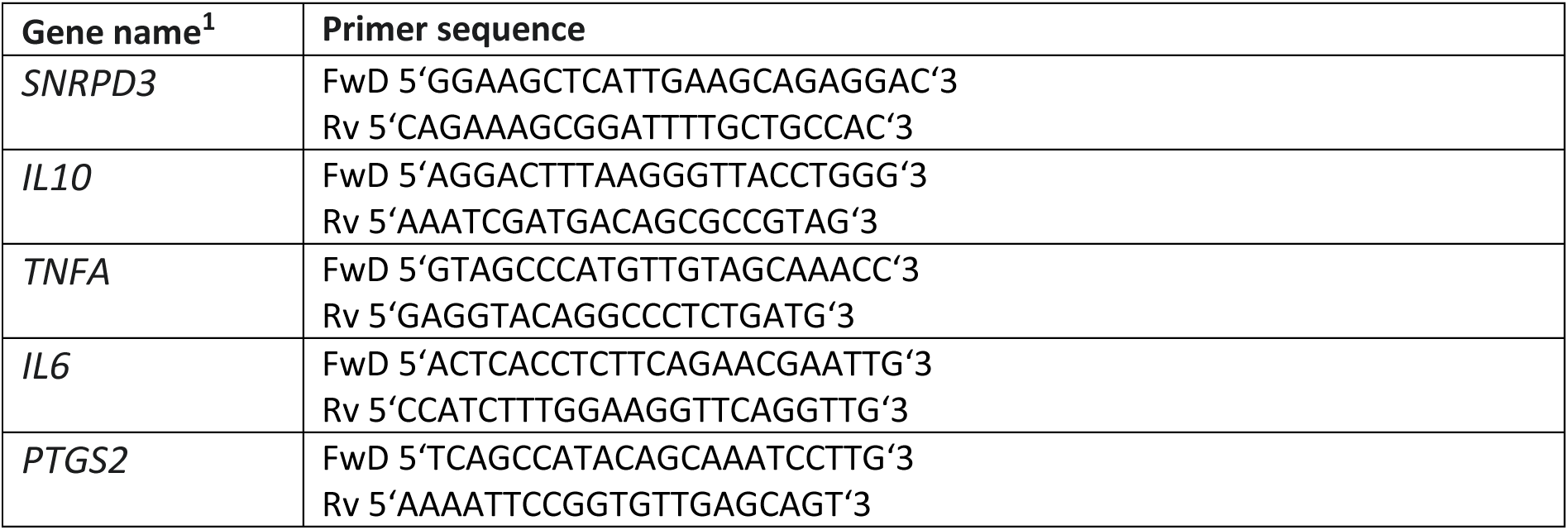
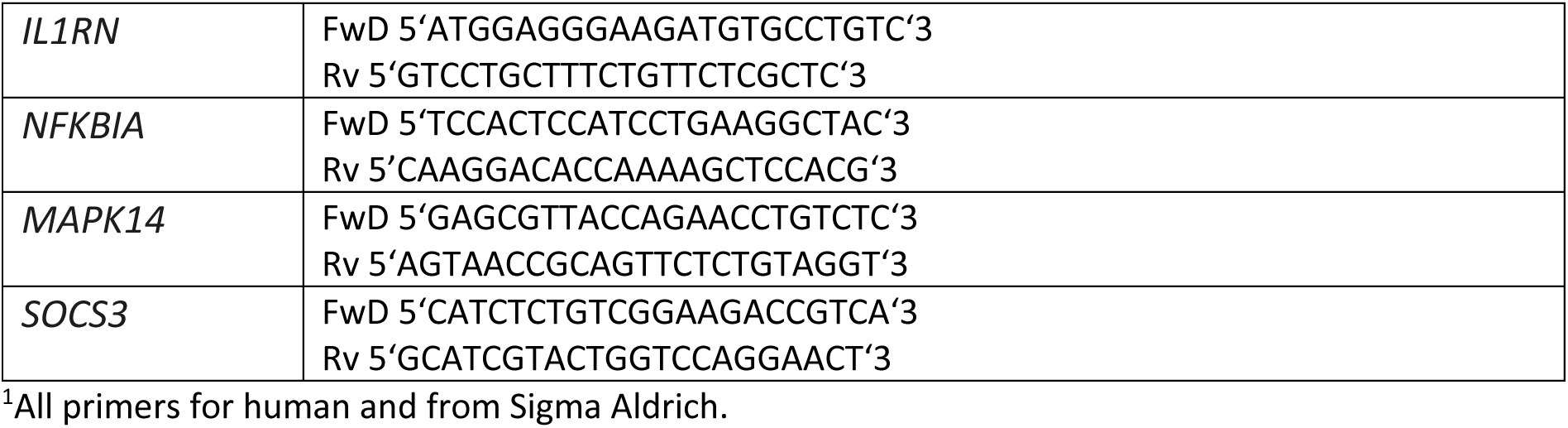
Primers for RT-qPCR.

### NF-κB reporter cell line

To investigate the effect of 4-HNE on NF-κB signaling, a monocytic NF-κB SEAP reporter cell line called THP-1 Blue NF-κB (Invivogen, thp-nfkbv2) was purchased. 100,000 cells per well were seeded in a 96-well U-bottom plate in RPMI 1640 (Gibco, 11530586), 10% heat-inactivated FBS (Hyclone, 10309433), 2 mM L-glutamine (Gibco, 15430614), 1% Antibiotic-Antimycotic (Gibco, 15240062) and incubated overnight at 37°C. The next day, cells were treated with different concentrations of 4-HNE in the presence and absence of LPS (100 ng/mL) (Sigma, L4391-1MG) for 6 hrs and 24 hrs, as described above. Thereafter, the assay was performed following manufacturer guidelines. In short, 180 μl QUANTI-blue solution (Invivogen, rep-qbs) was added to a 96-well flat bottom plate, and 20 μL of experimental supernatant was added on top before incubating at 37°C for 1 h. After incubation, the luciferase activity was measured with a Synergy Mx reader (BioTek) at OD of 655 nm.

### Western blot

For western blot, 1,000,000 macrophages were seeded in a 6-well plate. After experimental treatment, the cells were washed three times with cold PBS before lysing them with RIPA buffer (Thermo Scientific, 89900) containing Complete Mini EDTA-free Protease Inhibitor Cocktail (Sigma, 11697498001) and phosSTOP (Sigma/Merck, 4906837001) following manufacturer guidelines. Samples were denatured and unfolded using 4× Laemmli sample buffer (Bio-Rad, 1610747), heated at 90°C for 5 min, and separated using a 4–20% Mini-PROTEAN TGX Precast protein gels, 10 well, 50 μl (Bio-Rad, 456-1096) at 100 V. Proteins were blotted onto poly(vinylidene difluoride) (PVDF) membranes (Bio-Rad, 1620177) for 60 min at 90 V. Thereafter, blots were washed three times in UltraPure Water prior to blocking with Intercept Blocking Buffer (LiCor, 927-60001) for 60 min at room temperature. After blocking, blots were incubated with primary antibody (Table 4) overnight at 4°C. The next day, blots were washed three times and incubated for 60 min with secondary antibody at room temperature. Blots incubated with anti-rabbit IgG, HRP-linked Antibody were treated with WesternSure PREMIUM Chemiluminescent Substrate (LiCor, 926-95000). All membranes were imaged using the Li-Cor Odyssey Fc Imaging system and analyzed using Image Studio Software.

**Table 4:** Antibodies used for Western blot.

| Antibody | Company | Cat.: | Dilution |
| --- | --- | --- | --- |
| Phospho-p38 MAPK (Thr180/Tyr182) Antibody | Cell Signaling Technologies | 9211 | 1:1000 |
| p38 MAPK (D13E1) XP Rabbit mAb | Cell Signaling Technologies | 8690 | 1:1000 |
| Phospho-Stat3 (Tyr705) (D3A7) XP Rabbit mAb | Cell Signaling Technologies | 9145 | 1:500 |
| Stat3 (79D7) Rabbit mAb | Cell Signaling Technologies | 4904 | 1:500 |
| GAPDH (14C10) Rabbit mAb | Cell Signaling Technologies | 2118 | 1:2000 |
| GAPDH Antibody (G-9): sc-365062 | Santa Cruz | sc-365062 | 1:3000 |
| Donkey anti-rabbit IRDey 680 | LiCor | 926-32223 | 1:2000 |
| Donkey anti-mouse IRDey680 | LiCor | 926-32222 | 1:2000 |
| Anti-rabbit IgG, HRP-linked Antibody | Cell Signaling Technologies | 7074S | 1:1000 |

### MTT assay

Around 10,000 cells per well were seeded in a 96-well flat-bottom plate. Dimethyl sulfoxide (DMSO) (Sigma, 34869) treatment was used as a positive control. After treatment, cells were incubated with 0.5 mg/mL 3-(4,5-Dimethyl-2-thiazolyl)-2,5-diphenyl-2H-tetrazolium bromide (MTT) (Sigma Aldrich, M2128) solution for 3 hours at 37C. Subsequently, cells were washed once with PBS before incubating cells with DMSO for 10 min at 37C. The absorbance was measured at OD540 nm using a Synergy HTX multi-mode reader (BioTek).

### HIF-1α reporter assay

HRE-luciferase plasmid was a gift from Navdeep Chandel (*89*) (Addgene plasmid #26731), which was transfected into human monocyte-derived macrophages using the Neon transfection system (Fisher Scientific, 10090314) 10 μl tip protocol (1000V, 40 ms, 2 pulses). 2 μg of HRE-luciferase plasmid was used in T-buffer. Cells were cultured in an ultra-low adherence plate in phenol red-free RPMI (ThermoFisher, 11835030) containing 1% L-glutamine (Thermo Scientific, 15430614) and incubated for 1 hour at 37°C. Subsequently, 10% FBS and 100 ng/mL of M-CSF (R&D systems, 216-MC-500) were added for overnight recovery at 37°C.

The next day, cells were detached using cold PBS, and 100,000-150,000 cells were seeded in a 96-well U-bottom plate. Transfected cells were treated with 4-HNE in the presence or absence of LPS, as described above. Thereafter, firefly luciferase activity was determined using the Bright-Glo Luciferase assay system (Promega, E2610) following manufacture guidelines before measuring the luminescence using a Synergy Mx reader (BioTek).

### MitoSOX assay

Following manufacturer guidelines, the production of Mitochondrial ROS was measured using MitoSOX RED (ThermoFisher, M36008). In short, 100,000 cells were seeded in a 96-well U-bottom ultra-low adherence plate. During the last 30 minutes of the experiment, 1μM MitoSox was added before measuring the signal using a CytoFlex S flow cytometer (Beckman Coulter). Data were analysed using NovoExpress software (Agilent).

### Cell cytotoxicity by LDH release

5,000 cells/well were seeded in a 96-well U bottom plate. After the experiment, the supernatant was used to measure the LDH concentration using the CyQUANT LDH Cytotoxicity assay (ThermoFisher, C20300) according to manufacturer guidelines.

### Cytokine production by ELISA

The plasma from AcuteLines patients and the supernatant of 4-HNE-treated human monocyte-derived macrophages (see above) were used for ELISA to quantify the levels of TNF-α (ThermoFisher, 88-7346-88), IL-6 (ThermoFisher, 88-7066-88), and IL-10 (ThermoFisher, 88-7106). The ELISA was conducted according to manufacturer protocols.

### Statistics

Statistical analyses were performed using GraphPad Prism 10 software. The normal distribution was determined before data analysis. Data were tested with the Shapiro-Wilk normality test to check for a normal (Gaussian) distribution. Tests were used depending on the data distribution and the research question, as indicated in the figure legends. A p-value < 0.05 was considered significantly different.

### Ethics approval and consent to participate

This work has been carried out following The Code of Ethics of the World Medical Association (Declaration of Helsinki) for experiments involving humans. The Medical Ethics Review Committee (METc) of the UMCG reviewed and approved this study (METc; PaNaMa number 18642).

### Availability of data and materials

The datasets used and analyzed during the current study are deposited at the open-access repository Zenodo, and transcriptomics data have been deposited at the Gene Expression Omnibus. RNA sequencing data are deposited at the GEO Datasets, under accession: GSE289330.

## Acknowledgments

Acutelines provided the patient data and biomaterials used in this manuscript. Therefore, the authors would like to thank the Acutelines data and biobank, patients, their families, and healthy donors for contributing to this research.

## Supplementary materials

**Supplementary Figure 1:**
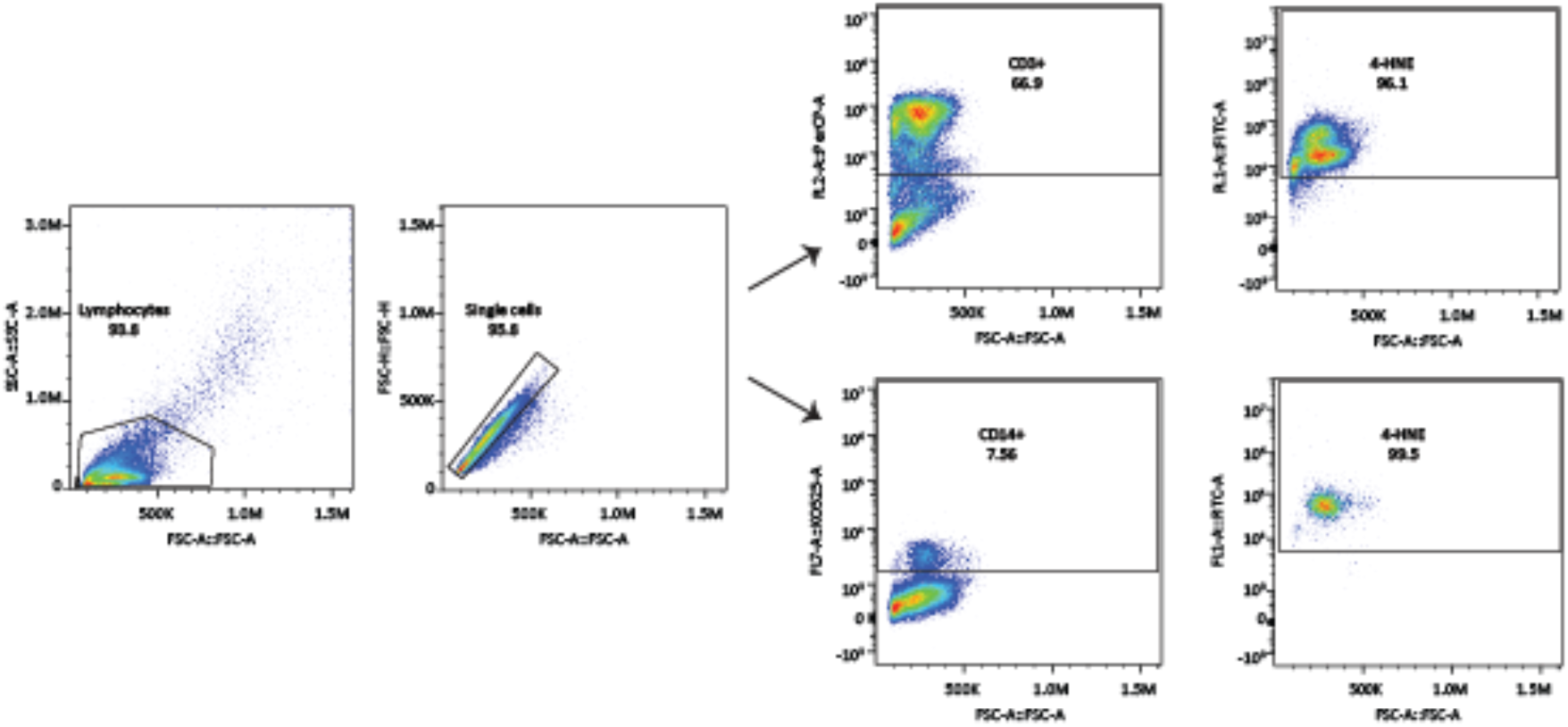
Gating strategy of Buffy coats stained with CD3^+^, CD14^+^, and 4-HNE. Buffy coats of Acuteline patients was stained for different cell surface markers and intracellular 4-HNE protein adducts. The gating strategy displayed is used for flow cytometry analysis.

**Supplementary Figure 2:**
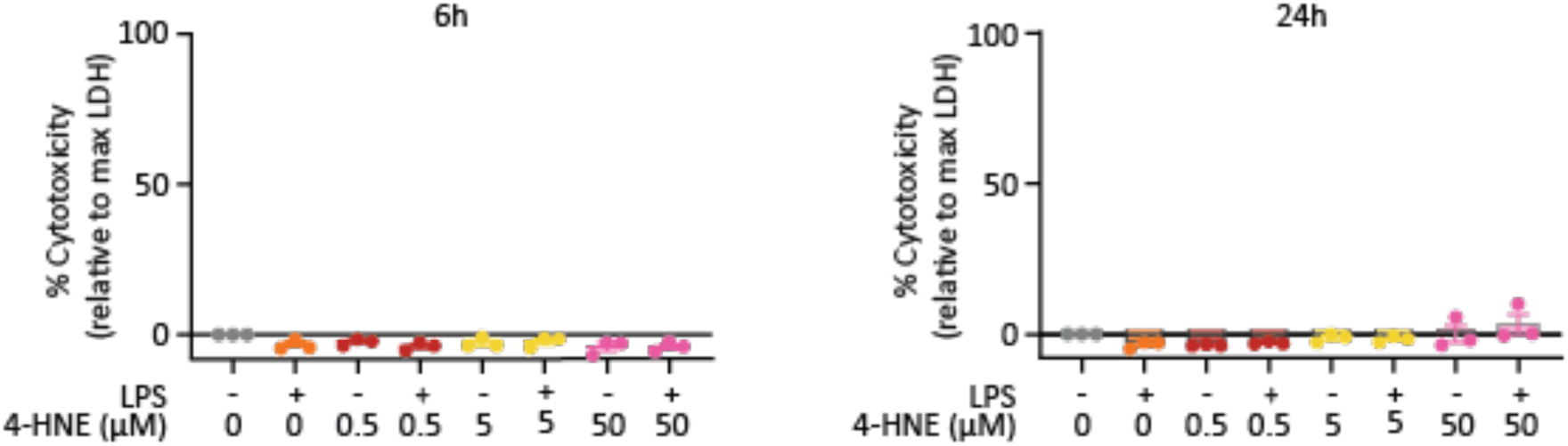
4-HNE treatment does not affect the viability of human macrophages. Macrophages were treated with and without 4-HNE in the presence or absence of LPS for 6h (left) and 24h (right). Cytotoxicity was determined relative to the maximal LDH production of 100% lysed cells. Each data point indicates one donor and the bar displays the SEM.

**Supplementary Figure 3:**
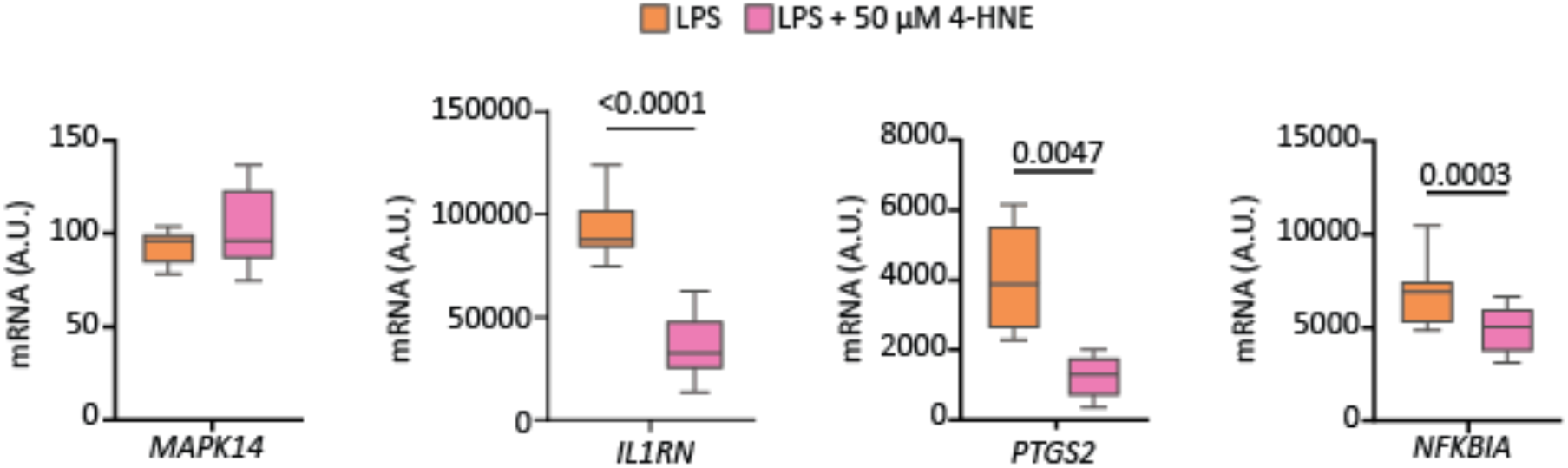
4-HNE treatment affects important immunoregulatory genes. Transcriptomics analysis of human monocyte-derived macrophages from 8 healthy donors treated with LPS (orange) or LPS + 50 μM 4-HNE (pink) for 24h. Data are displayed as arbitrary units of mRNA counts, and significance is tested with a paired t-test.

**Supplementary Figure 4:**
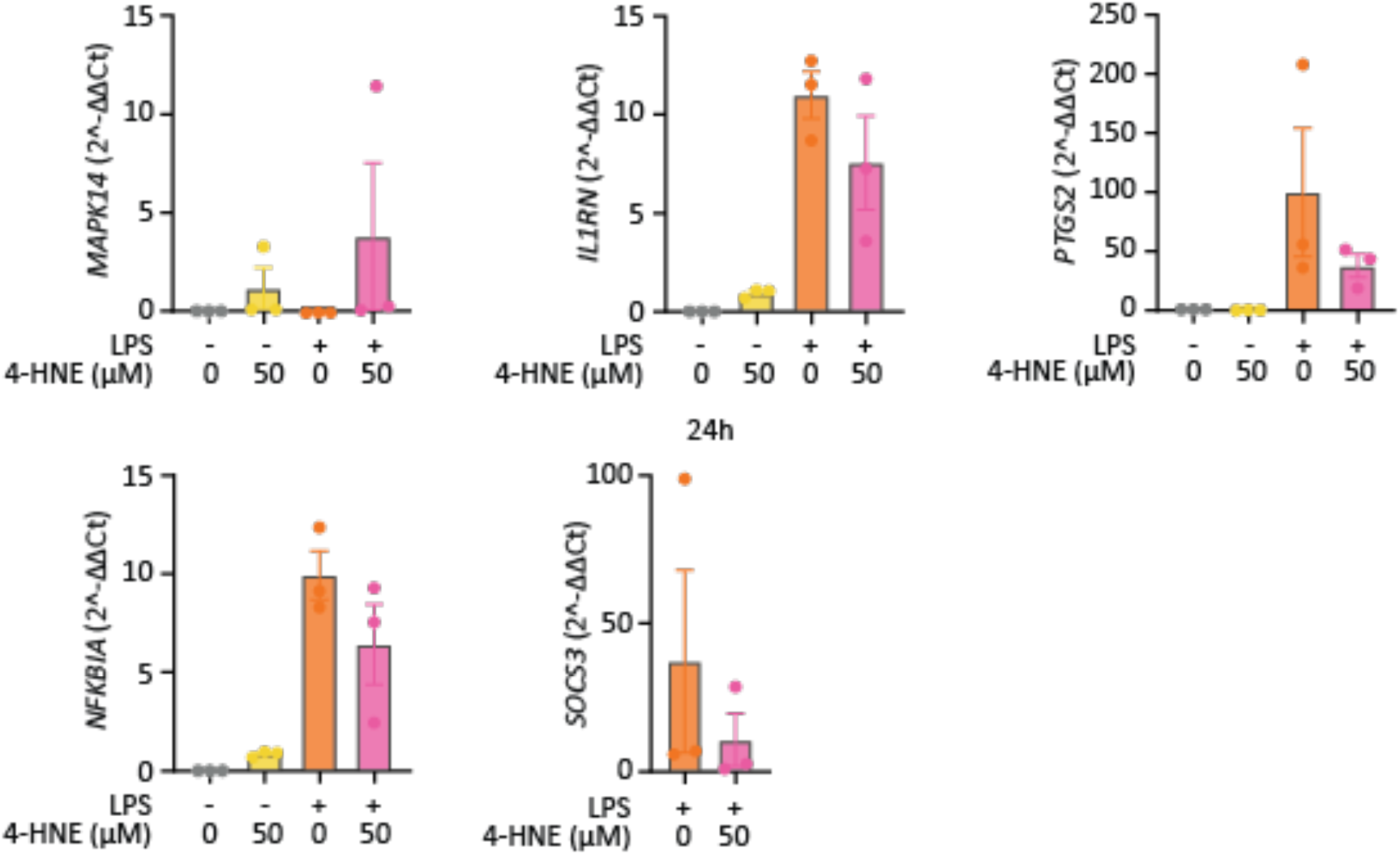
Quantification of the RNA sequencing data by RT-qPCR. Human monocyte-derived macrophages from 3 healthy donors were treated with LPS in the presence or absence of different concentrations of 4-HNE. Gene expression was measured after 24h by RT-qPCR. Each data point is one donor, and the error bar displays the SEM.

**Supplementary Figure 5:**
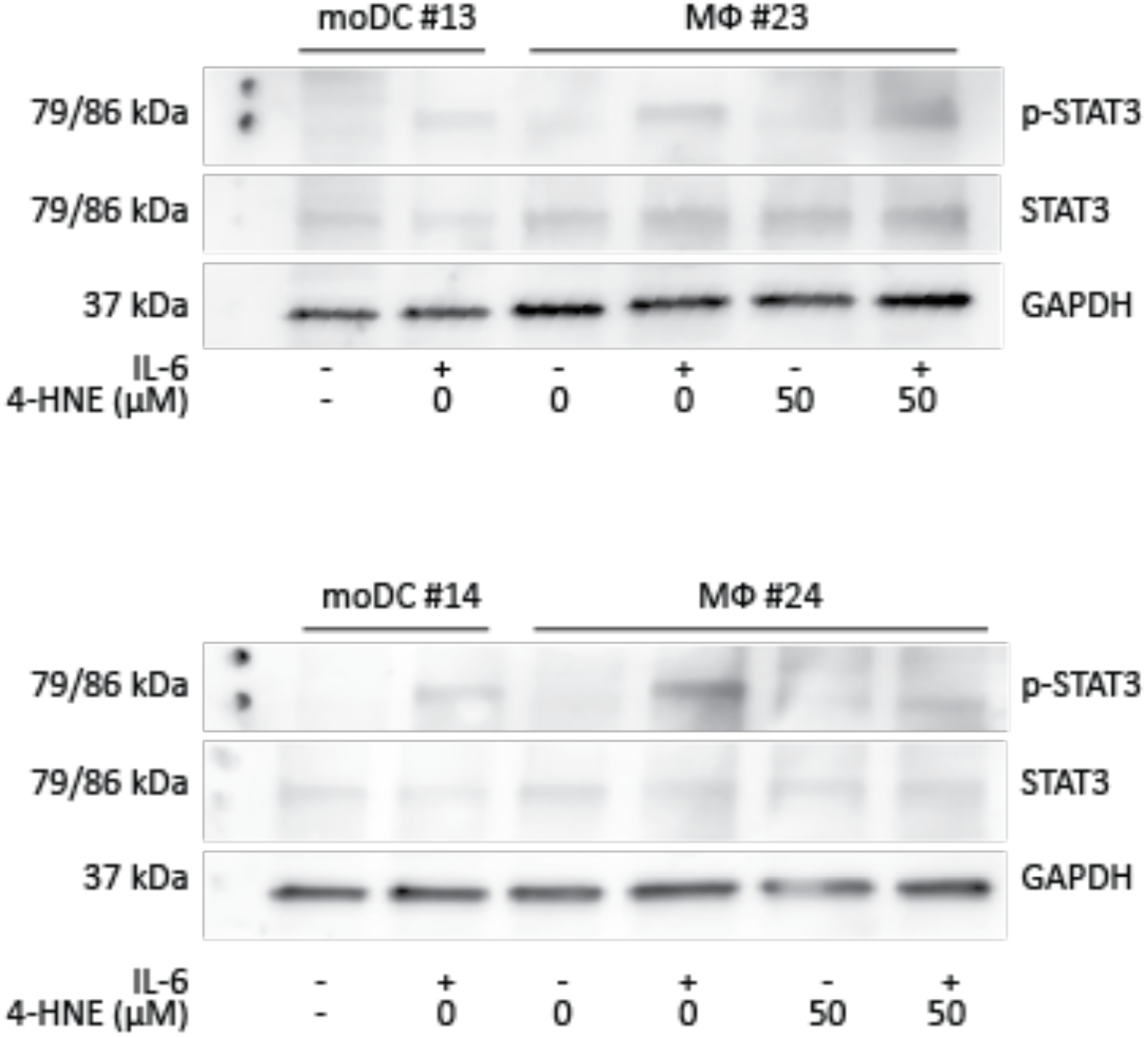
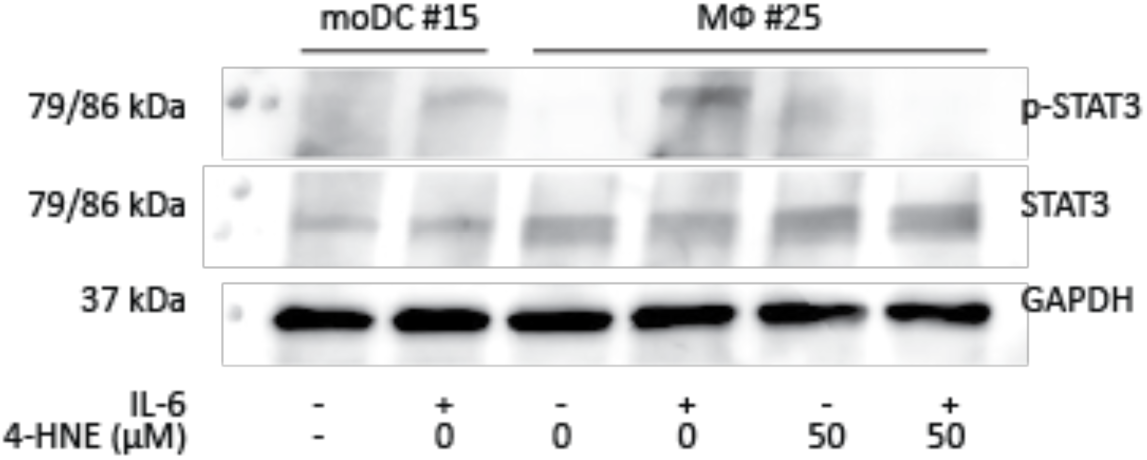
Western blot to determine STAT3 activity for three more donors. Western blots belonging to main figure 7B for monocyte-derived macrophages. Monocyte-derived dendritic cells (moDCs) treated with IL-6 are shown as a control.

**Supplementary Figure 6:**
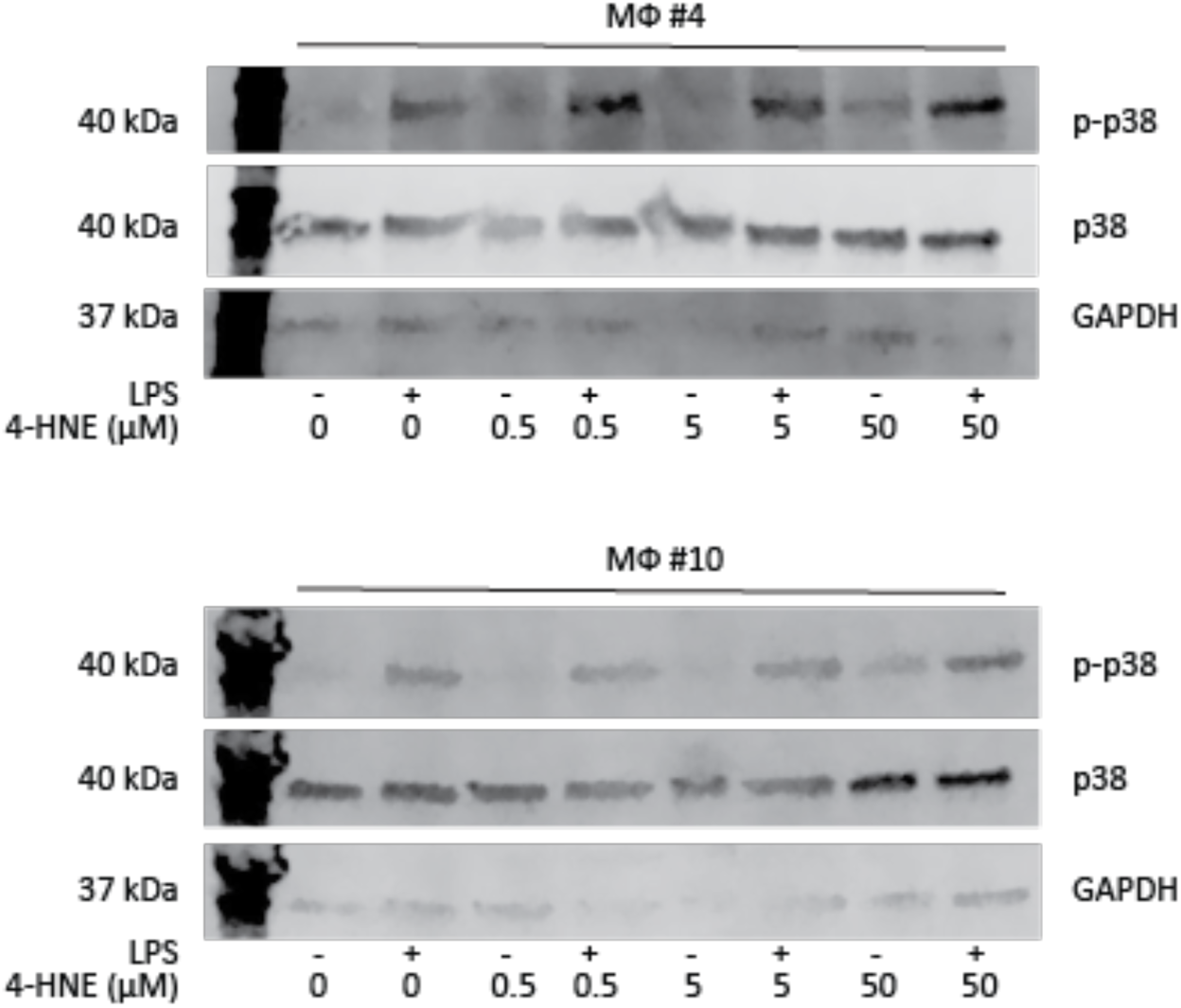
Western blot to determine p38 activity of second and third donors. Western blots belonging to main figure 9C.

**Supplementary Table 1:**
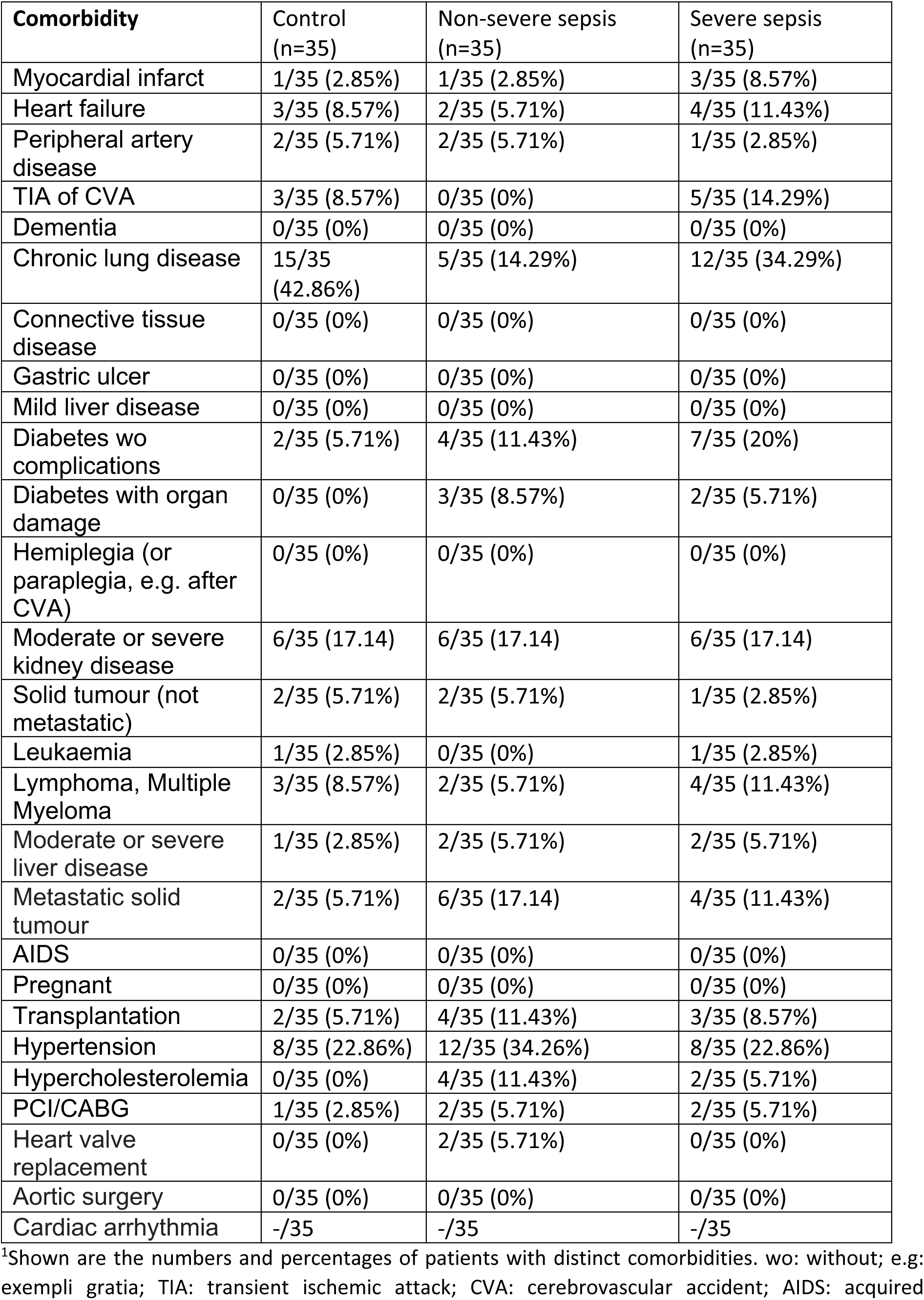
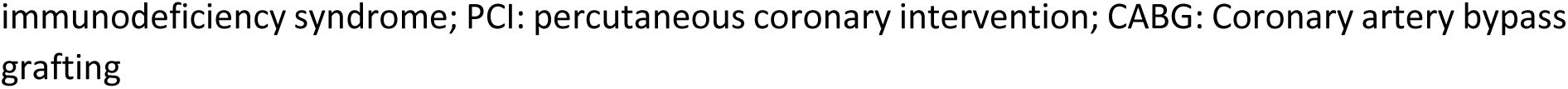
Comorbidities of patients from this study^1^.

| **Comorbidity** | **Control**<br>**(n=35)** | **Non-severe sepsis**<br>**(n=35)** | **Severe sepsis**<br>**(n=35)** |
| --- | --- | --- | --- |
| Myocardial infarct | 1/35 (2.85%) | 1/35 (2.85%) | 3/35 (8.57%) |
| Heart failure | 3/35 (8.57%) | 2/35 (5.71%) | 4/35 (11.43%) |
| Peripheral artery disease | 2/35 (5.71%) | 2/35 (5.71%) | 1/35 (2.85%) |
| TIA of CVA | 3/35 (8.57%) | 0/35 (0%) | 5/35 (14.29%) |
| Dementia | 0/35 (0%) | 0/35 (0%) | 0/35 (0%) |
| Chronic lung disease | 15/35 (42.86%) | 5/35 (14.29%) | 12/35 (34.29%) |
| Connective tissue disease | 0/35 (0%) | 0/35 (0%) | 0/35 (0%) |
| Gastric ulcer | 0/35 (0%) | 0/35 (0%) | 0/35 (0%) |
| Mild liver disease | 0/35 (0%) | 0/35 (0%) | 0/35 (0%) |
| Diabetes wo complications | 2/35 (5.71%) | 4/35 (11.43%) | 7/35 (20%) |
| Diabetes with organ damage | 0/35 (0%) | 3/35 (8.57%) | 2/35 (5.71%) |
| Hemiplegia (or paraplegia, e.g. after CVA) | 0/35 (0%) | 0/35 (0%) | 0/35 (0%) |
| Moderate or severe kidney disease | 6/35 (17.14) | 6/35 (17.14) | 6/35 (17.14) |
| Solid tumour (not metastatic) | 2/35 (5.71%) | 2/35 (5.71%) | 1/35 (2.85%) |
| Leukaemia | 1/35 (2.85%) | 0/35 (0%) | 1/35 (2.85%) |
| Lymphoma, Multiple Myeloma | 3/35 (8.57%) | 2/35 (5.71%) | 4/35 (11.43%) |
| Moderate or severe liver disease | 1/35 (2.85%) | 2/35 (5.71%) | 2/35 (5.71%) |
| Metastatic solid tumour | 2/35 (5.71%) | 6/35 (17.14) | 4/35 (11.43%) |
| AIDS | 0/35 (0%) | 0/35 (0%) | 0/35 (0%) |
| Pregnant | 0/35 (0%) | 0/35 (0%) | 0/35 (0%) |
| Transplantation | 2/35 (5.71%) | 4/35 (11.43%) | 3/35 (8.57%) |
| Hypertension | 8/35 (22.86%) | 12/35 (34.26%) | 8/35 (22.86%) |
| Hypercholesterolemia | 0/35 (0%) | 4/35 (11.43%) | 2/35 (5.71%) |
| PCI/CABG | 1/35 (2.85%) | 2/35 (5.71%) | 2/35 (5.71%) |
| Heart valve replacement | 0/35 (0%) | 2/35 (5.71%) | 0/35 (0%) |
| Aortic surgery | 0/35 (0%) | 0/35 (0%) | 0/35 (0%) |
| Cardiac arrhythmia | -/35 | -/35 | -/35 |

<sup>1</sup>Shown are the numbers and percentages of patients with distinct comorbidities. wo: without; e.g: exempli gratia; TIA: transient ischemic attack; CVA: cerebrovascular accident; AIDS: acquired
immunodeficiency syndrome; PCI: percutaneous coronary intervention; CABG: Coronary artery bypass grafting

